# Chromosomal domain formation by archaeal SMC, a roadblock protein, and DNA structure

**DOI:** 10.1101/2024.05.14.594075

**Authors:** Kodai Yamaura, Naomichi Takemata, Masashi Kariya, Sonoko Ishino, Masataka Yamauchi, Shoji Takada, Yoshizumi Ishino, Haruyuki Atomi

## Abstract

Structural maintenance of chromosomes (SMC) complexes fold genomes by extruding DNA loops. In eukaryotes, loop-extruding SMC complexes form topologically associating domains (TADs) by being stalled by roadblock proteins. It remains unclear whether a similar mechanism of domain formation exists in prokaryotes. Using high-resolution chromosome conformation capture sequencing, we show that an archaeal homolog of the bacterial Smc-ScpAB complex organizes the genome of *Thermococcus kodakarensis* into TAD-like domains. We also find that TrmBL2, a nucleoid-associated protein that forms a stiff nucleoprotein filament, stalls the *T. kodakarensis* SMC complex and establishes a boundary at the site-specific recombination site *dif*. TrmBL2 stalls the SMC complex at tens of additional non-boundary loci with lower efficiency. Intriguingly, the stalling efficiency is correlated with structural properties of underlying DNA sequences. Our study illuminates not only a eukaryotic-like mechanism of domain formation in archaea, but also an unforeseen role of intrinsic DNA structure in large-scale genome organization.

## INTRODUCTION

In both eukaryotes and prokaryotes, structural maintenance of chromosomes (SMC) complexes play critical roles in regulating the 3D structure and function of genomes. ^1–3^ At the core of the SMC complex is two SMC proteins featured by a ∼50-nm-long antiparallel coiled coil with the hinge dimerization domain at one end and the ATPase head domain at the other. The head domains of the dimer sandwich ATPs and hydrolyze them to drive conformational changes in the complex. The two SMC protomers are bridged by a Kleisin subunit to form a common tripartite ring structure. An intervening central domain of Kleisin further interacts with either Kleisin-interacting tandem winged-helix elements (Kites) or HEAT proteins associated with Kleisins (Hawks). ^4,5^

By employing single-molecule visualization and chromosome conformation capture (3C) techniques, researchers revealed that SMC complexes function as motors that progressively extrude DNA loops. ^6,7^ In eukaryotes, a major role of the SMC-mediated loop extrusion is to fold genomes into arrays of self-interacting domains. These structures, often called topologically associating domains (TADs) or loop domains, are believed to play versatile regulatory roles. ^6,8,9^ In most cases, eukaryotic chromosomal domains are formed by the SMC complex cohesin and a number of DNA-binding proteins (CTCF, RNA polymerases, etc.) that stall cohesin-mediated loop extrusion at domain boundaries. ^6,10–15^ Another eukaryotic SMC complex condensin also mediates domain formation in certain species, although it is less clear whether and how loop extrusion and boundary proteins are involved in this process. ^16,17^ As with eukaryotes, bacteria fold their genomes into arrays of self-interacting domains called chromosomal interaction domains (CIDs). ^18–20^ However, two major classes of bacterial SMC complexes, Smc-ScpAB and MukBEF, do not play a role in CID formation, and they instead use the loop extrusion activity to resolve replicated DNA molecules for chromosome segregation. ^18,20–27^ CIDs are formed by high levels of transcription occurring at their boundaries. ^18–20^ These findings have led to the prevailing view that domain formation driven by loop extrusion of SMC complexes is specific to eukaryotes.

Current evidence suggests the origin of eukaryotes within the prokaryotic domain Archaea. ^28,29^ Most archaea (with the notable exception of Crenarchaeota) possess homologs of the bacterial Smc-ScpAB subunits, namely the Smc ATPase, the Kleisin protein ScpA, and the Kite protein ScpB. ^30,31^ Recent *in vitro* experiments failed to detect a physical interaction between ScpA and ScpB from some archaea including *Pyrococcus yayanosii, Thermococcus onnurineus, Methanosalsum zhilinae* and *Methanothrix soehngenii*, despite the fact that their bacterial homologs form a stable subcomplex. ^30,32,33^ Genomes of certain archaeal lineages even lack *scpB* while they contain *smc* and *scpA*. ^30^ These findings led to the proposal that the archaeal homolog of Smc-ScpAB may function as a binary complex composed of only Smc and ScpA. However, this possibility has never been tested *in vivo*.

Our knowledge of archaeal 3D genome organization was considerably limited for a long time, partly due to a relatively low number of cultivated archaeal species, their small size, and the extreme growth conditions of most model archaea. ^34,35^ We and others have recently succeeded in applying genome-wide 3C techniques (Hi-C and 3C-seq) to diverse archaeal species, identifying a number of structural entities including self-interacting domains. ^36–42^ As is the case in bacteria, most of these archaeal chromosomal domains are demarcated by active transcription. ^36,38^ Intriguingly, formation of certain domain boundaries in the halophilic archaeon *Halofarax volcanii* depends on Smc rather than transcription. ^38^ This raises the possibility that SMC-mediated loop extrusion is a key driver of domain formation not only in eukaryotes but also in archaea. However, the underlying molecular mechanism is largely unknown, especially regarding how Smc-dependent boundaries are formed at specific loci.

To address this question, we set out to characterize the Smc-ScpAB homolog in the hyperthermophilic anaerobic archaeon *Thermococcus kodakarensis*. Like many other archaea, *T. kodakarensis* has a genome encoded in a single circular chromosome. Whereas *H. volcanii* lacks ScpB, *T. kodakarensis* possesses homologs of Smc (TK1017), ScpA (TK1018), and ScpB (TK1962). In the following, we provide evidence that these proteins act in concert with a roadblock nucleoid-associated protein (NAP) and sequence-dependent structural features of DNA to sculpt chromosomal domains.

## RESULTS

### Domain formation in the genome of *T. kodakarensis*

To investigate the role of Smc, ScpA, and ScpB for archaeal genome organization, we developed a high-resolution 3C-seq protocol for *T. kodakarensis*, starting from the Hi-C procedure for prokaryotes published recently. ^38^ In this 3C-seq protocol, we used two crosslinking reagents (formaldehyde and disuccinimidyl glutarate) to capture fine-scale structures. ^43^ Crosslinked DNA was digested with two four-base blunt-end cutters (AluI and HaeIII) for high resolution and uniform digestion. We applied the method to our laboratory strain KU216, a uracil-auxotrophic strain (*ΔpyrF*) constructed from the wild-type *T. kodakarensis* strain KOD1. ^44^ Cells were grown in nutrient-rich medium (ASW-YT-m1- S^0^) until mid-to-late log phase and immediately fixed for the experiment. An obtained contact map, binned at 5-kb resolution, was largely similar to that of KOD1 published previously(Figures 1A and S1A). ^38^ Our 3C-seq data also generated a high-coverage contact map even at 500-bp resolution, whereas the published Hi-C data generated a highly sparse contact map at this resolution (Figures 1B and S1B). Since previous Hi-C/3C-seq analyses of archaea were conducted at resolutions ranging from 1 to 30 kb, ^36–42^ this study has provided the highest-resolution view of archaeal genome conformation.

**Figure 1.**
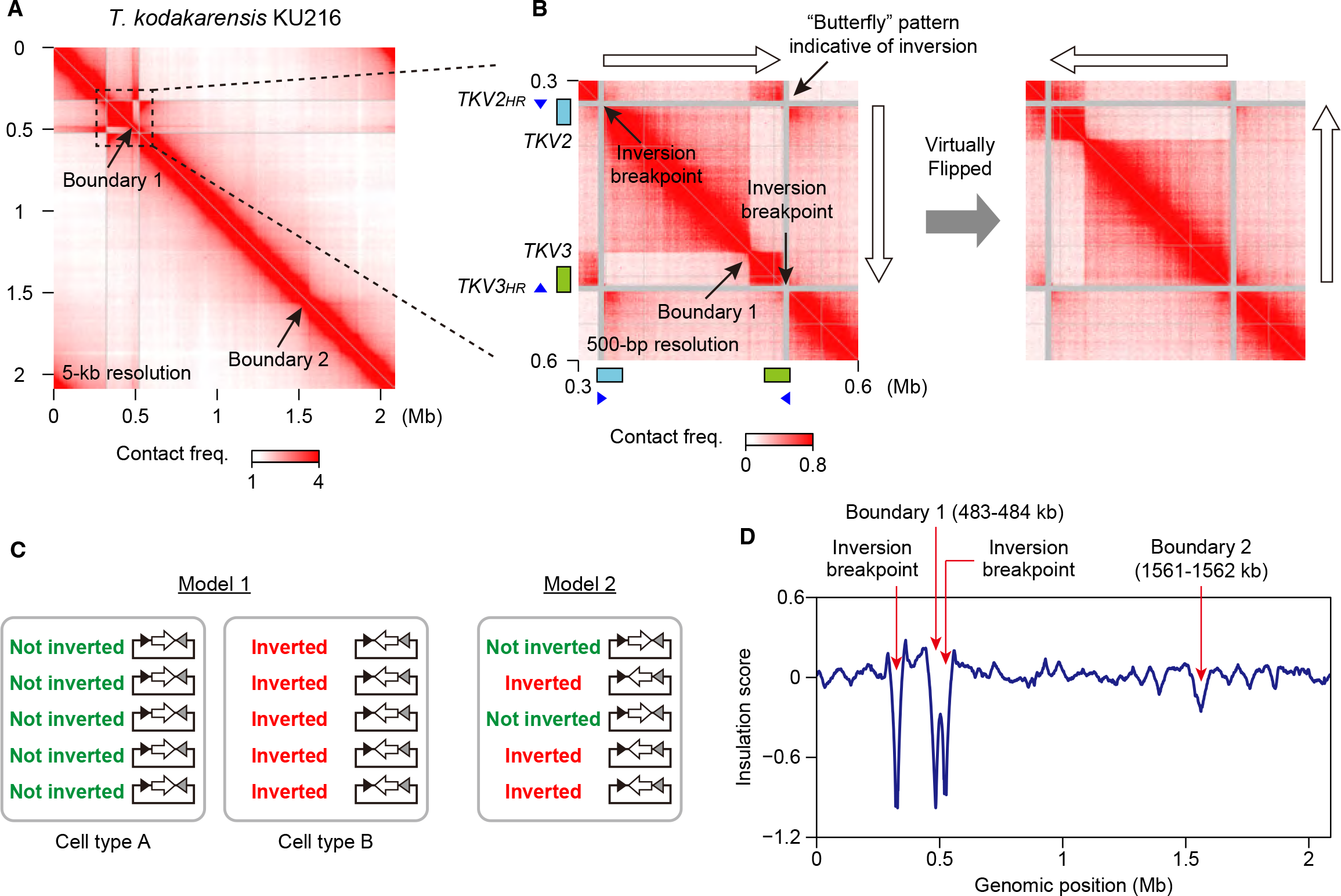
Identification of a genomic inversion and chromosomal domains in *T. kodakarensis* (A) 3C-seq analysis on the genome of the *T. kodakarensis* strain KU216. The contact map was generated at 5-kb resolution. Two visible boundaries (boundary 1 and boundary 2) are indicated by arrows. (B) Left panel: magnified 3C-seq contact map (500-bp resolution) showing a butterfly pattern indicative of a genomic inversion. Two proviral regions (*TKV2* and *TKV3*) are indicated by cyan and green rectangles, respectively. Homologous sequences within *TKV2* and *TKV3* (*TKV2HR* and *TKV3HR*, respectively) are indicated by blue triangles. Right panel: a 3C-seq contact map was generated as in the left panel, except that it was generated for a virtual genome sequence in which a DNA segment flanked by the expected inversion breakpoints was flipped. (C) Models for the suggested co-existence of chromosome copies with and without the inversion. (D) Insulation score profile of the KU216 genome. Note that the genomic inversion has caused an artificial drop in the contact insulation at the breakpoints, manifested as deep valleys in the profile.

Of note, the 3C-seq contact map of KU216 displayed a butterfly-like pattern indicative of genomic inversion (Figure 1B, left panel). ^45^ Expected inversion breakpoints were located in highly homologous ∼9-kb sequences (sequence identity: 96%) in the two proviral regions *TKV2* and *TKV3*.

^46^ These sequences, here denoted as *TKV2HR* and *TKV3HR* respectively, are oriented in opposite directions on the chromosome, which probably caused the inversion via intra-chromosomal crossover. As *T. kodakarensis* contains 7-19 copies of the chromosome per cell, ^47^ we wondered whether the inversion exists in all copies of the chromosome in the KU216 cell. To test this, we re-generated a 3C-seq contact map of KU216 using a virtual reference genome sequence in which the intervening segment between *TKV2HR* and *TKV3HR* was inverted. This manipulation did not eliminate the butterfly pattern (Figure 1B, right panel), suggesting that chromosome copies with and without the inversion co-exist at the single-cell level or the population level (Figure 1C). Published Hi-C data of the wild-type strain KOD1^38^ also generated a weaker but visible butterfly signal on the original reference genome (Figure S1B). Thus, the genomic heterogeneity is not specific to our laboratory strain, and chromosome copies harboring the inversion can exist at a variable ratio.

The 3C-seq contact map of KU216 displayed two visible boundaries across which genomic contacts were relatively depleted (Figure 1A). These boundaries, denoted as boundary 1 and boundary 2 respectively, can also be seen in the published Hi-C contact map of KOD1 (Figure S1A). ^38^ To determine the precise locations of the boundaries, we used the metric called insulation score, ^17^ which represents the relative frequency of local contacts across a locus and thereby reflects contact insulation strength of the region. By determining local minima of insulation scores at 1-kb resolution, we determined the positions of boundaries 1 and 2 as 483-484 kb and 1561-1562 kb, respectively (Figure 1D). The insulation of local contacts was much weaker at other loci, suggesting that boundaries 1 and 2 are the major boundaries on the *T. kodakarensis* genome.

The euryarchaeon *H. volcanii* and members of the Crenarchaeota form tens of DNA loops. ^36,38,39^ To search for DNA loops in *T. kodakarensis*, we analyzed the 3C-seq data of KU216 using Chromosight. ^48^ This analysis detected ∼10 loops from each of three replicates, but none of them were reproducible in all the replicates (data not shown). Loop anchors might be obscured by the combined effect of the polyploidy and thermal motion of the chromosome under the high growth temperature of *T. kodakarensis*.

### Smc, ScpA, and ScpB are all required to form chromosomal domains

To investigate whether Smc, ScpA, and ScpB contribute to boundary formation in *T. kodakarensis*, we constructed five deletion mutants lacking one or two of these proteins (Δ*smc*, Δ*scpA*, Δ*scpB*, Δ*smc* Δ*scpA*, and Δ*smc* Δ*scpB*) using KU216 as a parental strain. 3C-seq uncovered that all deletions tested slightly increased short-range interactions up to ∼200 kb (Figures S2A and B). More importantly, all deletions resulted in loss of contact insulation at boundaries 1 and 2, especially at the former (Figures 2A and B). The disruption of these boundaries was also manifested as an increase in the insulation score for both loci (Figure 2C). The score was increased very similarly among the five mutants. From these results, we conclude that Smc, ScpA, and ScpB act in the same pathway to organize the *T. kodakarensis* genome into the domain structures. Our unpublished results also suggest that these three proteins form a ternary complex (see Discussion). According to these findings, we will refer to the *T. kodakarensis* counterpart of bacterial Smc-ScpAB simply as Smc-ScpAB.

**Figure 2.**
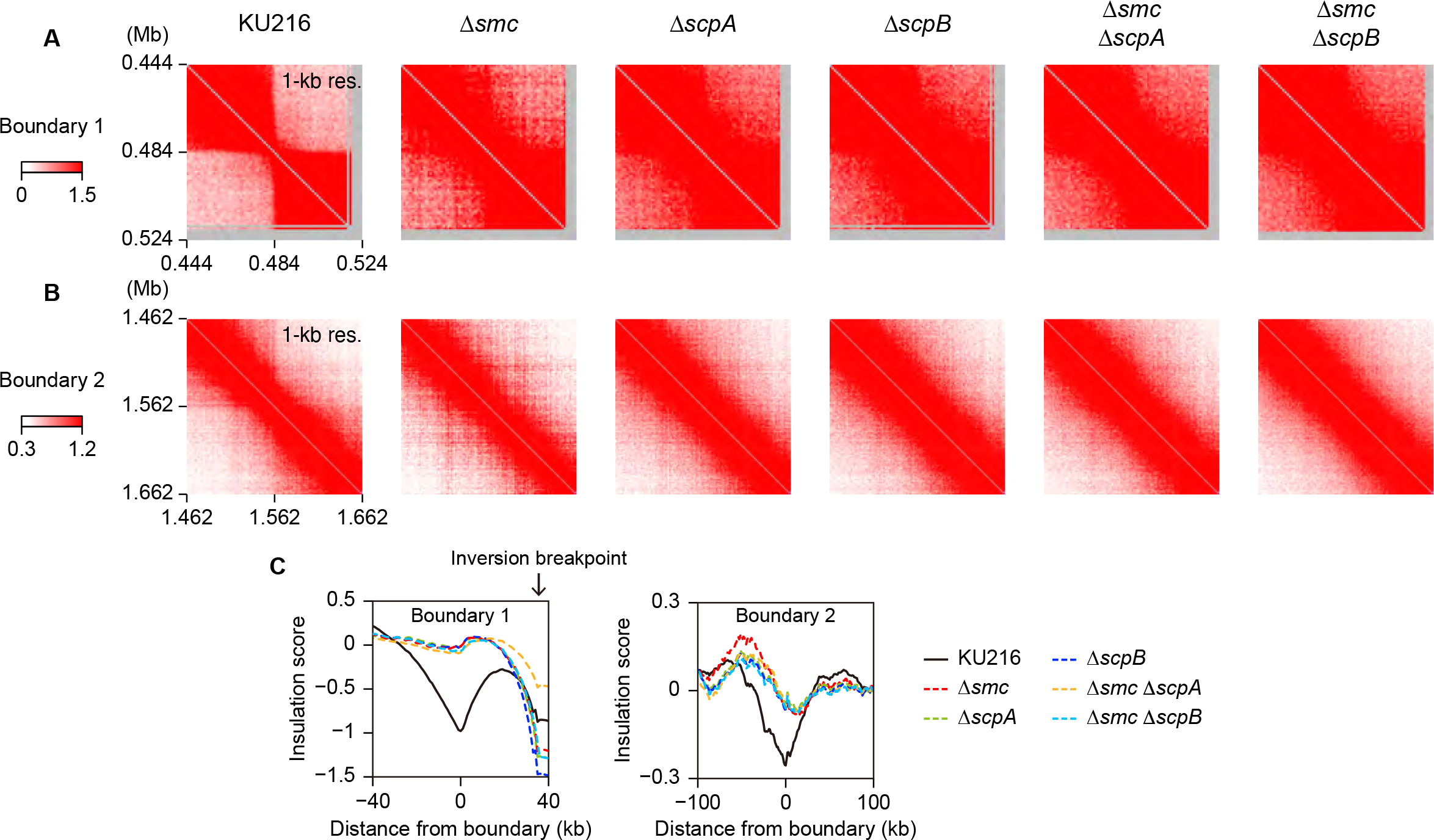
Deletions of Smc, ScpA, and ScpB decrease contact insulation at boundaries 1 and 2 in a non-additive manner (A and B) 3C-seq contact maps of genomic regions surrounding boundary 1 (A) and boundary 2 (B). KU216 and derivative strains lacking one or two of the Smc-ScpAB subunit homologs were used for the analysis. (C) Insulation score profiles of boundary 1 (left panel) and boundary 2 (right panel) are shown for KU216 and the deletion strains. Note that the graphs in the left panel cover an inversion breakpoint that has caused artificial drops in the score.

SMC-mediated genome organization is critical for cell viability in the bacterial model organisms *Escherichia coli* and *Bacillus subtilis*. ^21,22^ An early study reported that loss of *smc* causes a discernible growth defect in the archaeon *Methanococcus voltae*, whereas Cockram et al. more recently revealed that deletion of *smc* has no apparent impact on the cellular fitness of *H. volcanii*. ^38,49^ To explore whether the SMC-mediated domain formation has phenotypic consequences in *T. kodakarensis*, we first measured the growth of our deletion mutants in nutrient-rich medium. All of them proliferated at similar rates as the parental strain during exponential phase, although the mutants exhibited a slight decrease in growth rate immediately before the transition to stationary phase (Figure S3A). We also carried out RNA-seq analysis using the Δ*smc*, Δ*scpA*, and Δ*scpB* strains grown to mid- to-late exponential phase, observing only minor differences in their transcriptomes compared to that in the parental strain (Figure S3B). These results suggest that the domain structures shaped by Smc- ScpAB have a very small impact on the cellular fitness of *T. kodakarensis*, at least under the growth conditions tested here.

### Boundary formation near *dif* sequences in diverse euryarchaea

Boundary 1 was located in a large intergenic region (∼1.9 kb) between the convergently oriented genes *TK0561* and *TK0562*. We found that this intergenic region also contains a previously reported putative *dif* sequence composed of imperfect inverted repeats of 11 bp separated by a 6-bp spacer (Figures 3A and S4A). ^50^ *dif* is known as a target site of Xer site-specific recombinases (XerC and XerD in bacteria and XerA or Xer in archaea), which resolve chromosome dimers arising from an odd number of crossover events between circular sister chromosomes. ^51,52^ Chromatin immunoprecipitation sequencing (ChIP-seq) on the sole homolog of XerA (TK0777) in *T. kodakarensis* showed that the *dif* sequence was the *bona fide* binding site for XerA (Figure 3B). We did not find any discernible DNA motif around boundary 2.

**Figure 3.**
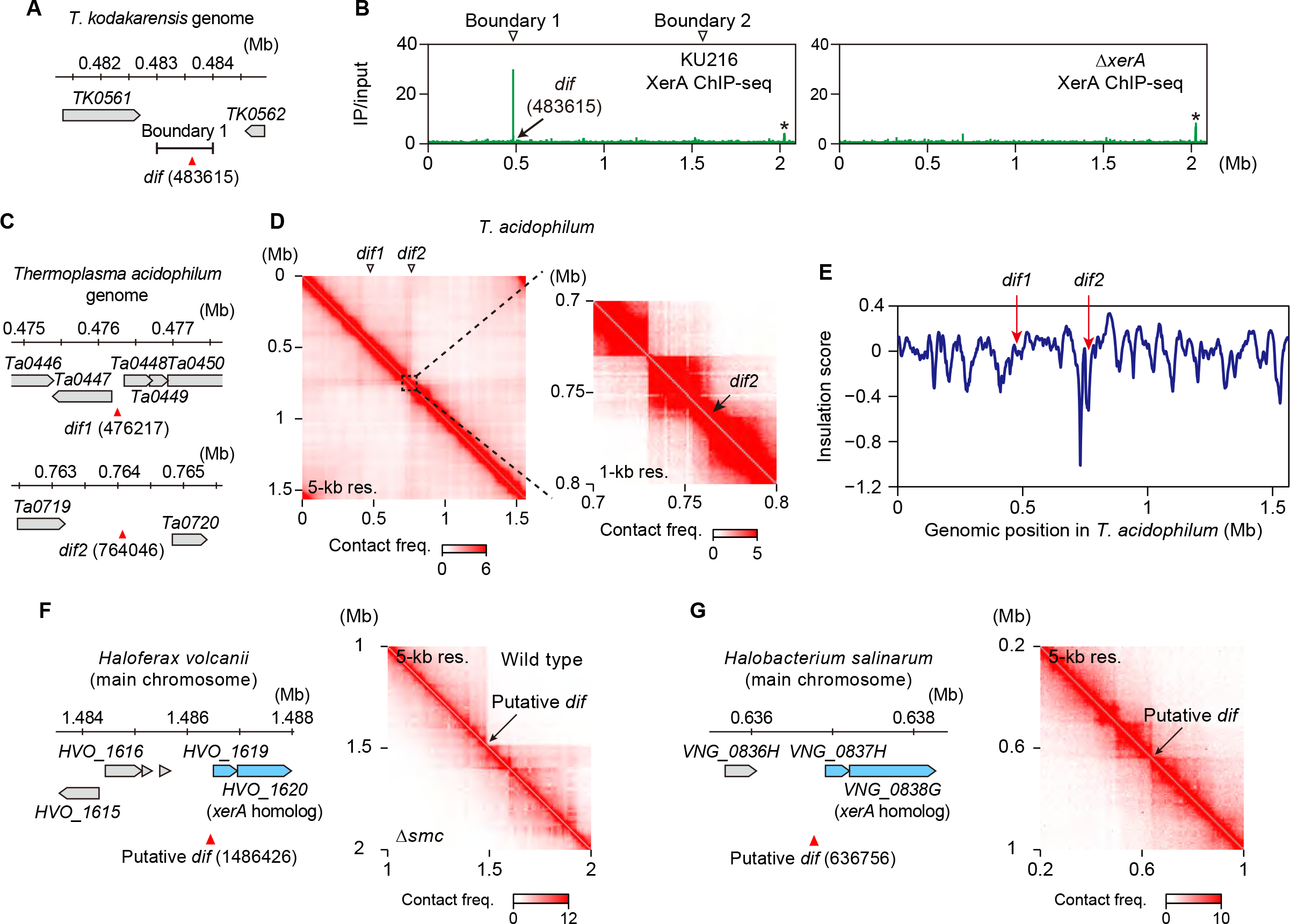
Domain boundaries are formed near *dif* sequences in diverse euryarchaea (A) Genomic position of the *dif* sequence in *T. kodakarensis*. Coordinates of the first bases of the *dif* sequences are in parentheses. The position of boundary 1 (determined at 1-kb resolution) is also shown. Neighboring genes and their orientations are indicated by gray pentagons. (B) ChIP-seq profiles for XerA in KU216 and Δ*xerA* cells of *T. kodakarensis*. Locations of boundaries 1 and 2 are indicated by white triangles. The *dif* position is also indicated by an arrow. The coordinate of the first base of the *dif* sequences is in parentheses. An asterisk indicates a non-specific peak. (C) Genomic positions of *dif1* and *dif2* in *Thermoplasma acidophilum*^96^ are shown as in (A). (D) Left panel: 3C-seq analysis on the genome of *T. acidophilum*. The contact map was generated at 5-kb resolution. Right panel: magnified contact map (1-kb resolution) showing the colocalization of *dif2* with a boundary structure. (E) Insulation score profile of the *T. acidophilum* genome. The genomic positions of *dif1* and *dif2* are indicated by red arrows. (F) Left panel: the genomic position of a putative *dif* sequence on the main chromosome of *Haloferax volcanii*. The coordinate of the first base of the *dif* sequence is in parentheses. Neighboring genes and their orientations are indicated by gray and cyan pentagons. Highlighted in cyan is an operon containing a homolog of *xerA*. Right panel: published Hi-C data^38^ were used to generate contact maps (5-kb resolution) around the *dif* sequence in wild-type and Δ*smc* cells of *H. volcanii* (upper right and lower left triangles, respectively). (G) Left panel: the genomic position of a putative *dif* sequence is shown as in (E) for the main chromosome of *Halobacterium salinarum*. Right panel: published Hi-C data^38^ were used to generate a contact map (5-kb resolution) around the *dif* sequence in wild-type *H. salinarum*.

A previous study identified two XerA-binding sites (*dif1* and *dif2*) in *Thermoplasma acidophilum*, a euryarchaeon possessing homologs of Smc, ScpA, and ScpB (Figure 3C). Of these two sites, only *dif2* can serve as a substrate for XerA-mediated recombination *in vitro*. ^53^ By applying 3C-seq to *T. acidophilum*, we identified a boundary-like structure near *dif2* but not *dif1* (Figures 3D and E). To further explore the generality of *dif*-associated boundaries, we searched for *dif*-like sequences around the previously reported Smc-dependent boundaries in the multipartite genome of *H. volcanii*. ^38^ We found that two of the nine Smc-dependent boundaries, one of which resides on the main chromosome and the other on the megaplasmid pHV3, are adjacent to putative *dif* sequences (Figures 3F and S4B). The *dif*-like element on the main chromosome is located upstream of an operon containing a *xerA* homolog (Figure 3F, left panel). We also inspected a published Hi-C dataset of *Halobacterium salinarum*, another euryarchaeon with a multipartite genome. ^38^ In this organism, a *dif*-like sequence is found upstream of the same *xerA-*containing operon and located close to a boundary structure (Figure 3G). It remains unknown whether this boundary is formed by an SMC complex. ^38^ Taken together, the colocalization of a boundary structure and *dif* is conserved among a wide range of euryarchaeal lineages.

A recent study has shown that bacterial XerD also serves as an unloader of Smc-ScpAB at the *dif*-like sequences named *XDS* in the replication terminus. ^54^ Presumably due to this unloading function, an artificial array of *XDS* inserted on a bacterial chromosome inhibits the translocation of Smc-ScpAB and forces the complex to form a boundary-like structure. ^54^ These findings led us to hypothesize that archaeal XerA is responsible for specifying the *dif*-associated boundary. To test this possibility, we tried to construct *T. kodakarensis* strains lacking either XerA or the whole 28-bp sequence of *dif* using KU216 as a parental strain. Consistent with the successful deletion of Xer in the crenarchaeon *Saccharolobus solfataricus*, ^51^ we obtained both mutants of *T. kodakarensis*. To our surprise, 3C-seq revealed that the formation of boundary 1 was completely normal in these mutants (Figures 4A and B). In addition, conformational changes in other genomic regions were not observed in Δ*xerA* or Δ*dif* (Figures S4C and D). Altogether, despite the conserved colocalization of a boundary and archaeal *dif*, the Xer/*dif* system is not essential for the boundary formation in *T. kodakarensis*.

**Figure 4.**
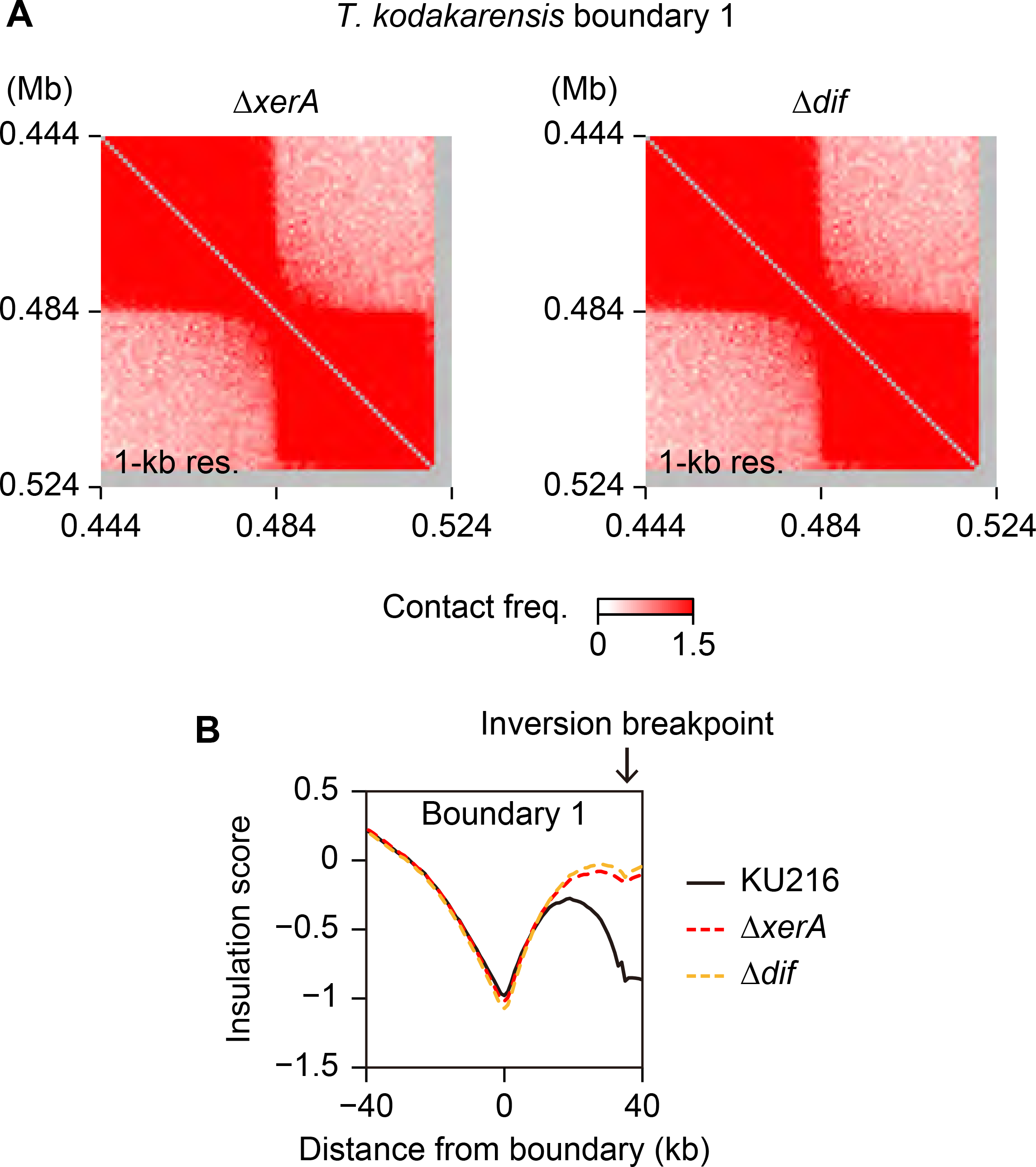
The Xer/*dif* system is dispensable for the formation of boundary 1 (A) 3C-seq contact maps (1-kb resolution) of a genomic region surrounding boundary 1 are shown for *T. kodakarensis* strains lacking either *xerA* (left panel) or *dif* (right panel). (B) Insulation score profiles of boundary 1 from KU216, Δ*xerA*, and Δ*dif* strains of *T. kodakarensis*. Note that the graphs cover an inversion breakpoint that has caused artificial drops in the score.

### The NAP TrmBL2 is required for the formation of boundary 1

To identify other candidates that specify boundary 1, we performed DNA affinity purification of proteins that bind to the intergenic region encompassing boundary 1. We prepared two biotinylated DNA probes, denoted as *difL* and *difR* respectively, that cover the intergenic region (Figure 5A). The kanamycin resistance gene *kanR* was used as a control probe. These probes were conjugated to streptavidin beads and incubated with cell extracts of KU216 in the presence of 150 mM NaCl. After washing with 150 mM NaCl, bound proteins were eluted in a single step by adding SDS (“total bound proteins” in Figure 5B). The *difL* and *difR* probes yielded two common specific bands (bands 1 and 2) as well as many other non-specific bands (Figure 5C). We reduced these non-specific proteins by eluting the bound material with increasing concentrations of NaCl before adding SDS. Bands 1 and 2 were isolated from two eluates (eluates 1 and 2) and analyzed by mass spectrometry (Figures 5B and C). Band 1 was identified as TrmBL2 (TK0471), a NAP known to form a stiff nucleoprotein filament and repress more than a hundred of genes. ^55,56^ Band 2 was identified as a putative homolog of the modified cytosine restriction protein McrB (TK0795). ^57^ Although both *difL* and *difR* contained *dif*, we did not see a clear specific band corresponding to XerA (33 kDa) under our experimental conditions.

**Figure 5.**
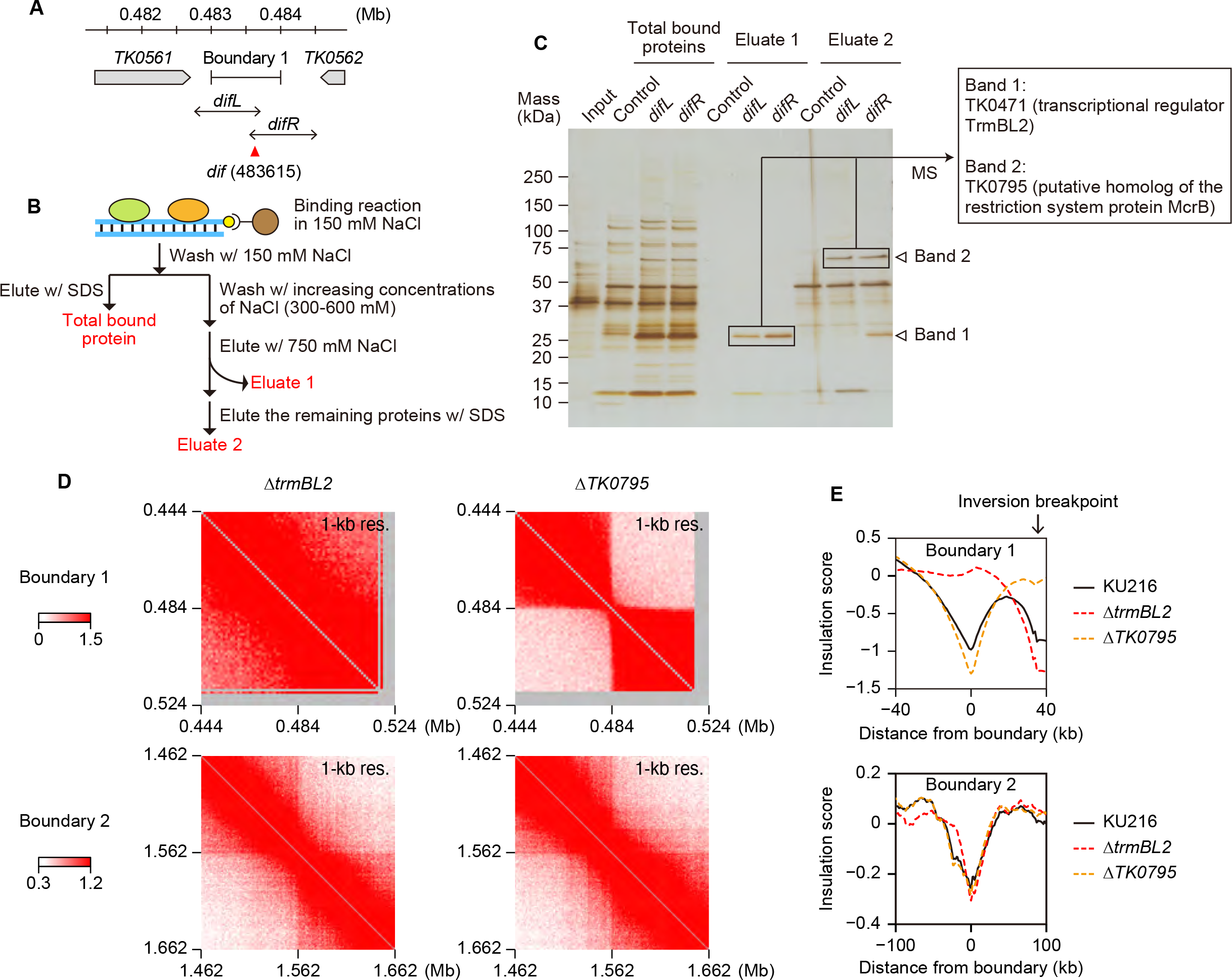
Identification of boundary 1-associated proteins and their impacts on genome architecture (A) DNA probes used for DNA affinity purification. (B) Workflow of DNA affinity purification. (C) Proteins from the DNA affinity purification experiment were separated by SDS-PAGE and visualized by silver staining. Positions of common specific bands for the *difR* and *difL* probes are indicated by white triangles (bands 1 and 2). Bands indicated by black rectangles were subjected for mass spectrometry. Input corresponds to 0.1% of the lysate used for each purification. (D) 3C-seq contact maps of genomic regions surrounding boundary 1 (top panels) and boundary 2 (bottom panels). KU216-derived strains lacking either *trmBL2* or *TK0795* were used for the analysis. (E) Insulation score profiles of boundary 1 (top panel) and boundary 2 (bottom panel) are shown for KU216 and deletion strains. Note that the graphs on the top panel cover an inversion breakpoint that has caused artificial drops in the score.

To investigate whether the identified proteins are required to build boundary 1, we constructed deletion mutants of *trmBL2* and *TK0795* using KU216 as a parental strain. 3C-seq demonstrated that the *trmBL2* deletion caused a complete loss of boundary 1, while the *TK0795* deletion rather slightly enhanced the contact insulation at boundary 1 (Figures 5D and E). The insulation of contacts was largely maintained at boundary 2 in both mutants. Except for the loss of boundary 1 in Δ*trmBL2*, we did not see a clear structural change in either of the mutants (Figure S5). Taken together, TrmBL2 is responsible for specifying boundary 1, while another, unknown factor defines boundary 2.

### TrmBL2 stalls Smc-ScpAB at boundary 1 and tens of other loci

We wondered whether TrmBL2 establishes a boundary by stalling loop extrusion as seen for eukaryotic CTCF. ^6,10^ To test this possibility, we carried out ChIP-seq of Smc and TrmBL2 using antisera against these proteins. The specificity of the antisera was confirmed by performing ChIP-seq on Δ*smc* and Δ*trmBL2* cells (Figures S6A and B). On the genome of KU216, Smc was most highly enriched at boundary 1 (Figure S6A). At a finer scale, boundary 1 was nestled by two sharp peaks of Smc, each of which overlapped with a distinct TrmBL2 peak (Figure 6A). Furthermore, these Smc peaks were almost completely lost in Δ*trmBL2* cells. These results support the idea that TrmBL2 forms boundary 1 by stalling Smc-ScpAB at this locus.

**Figure 6.**
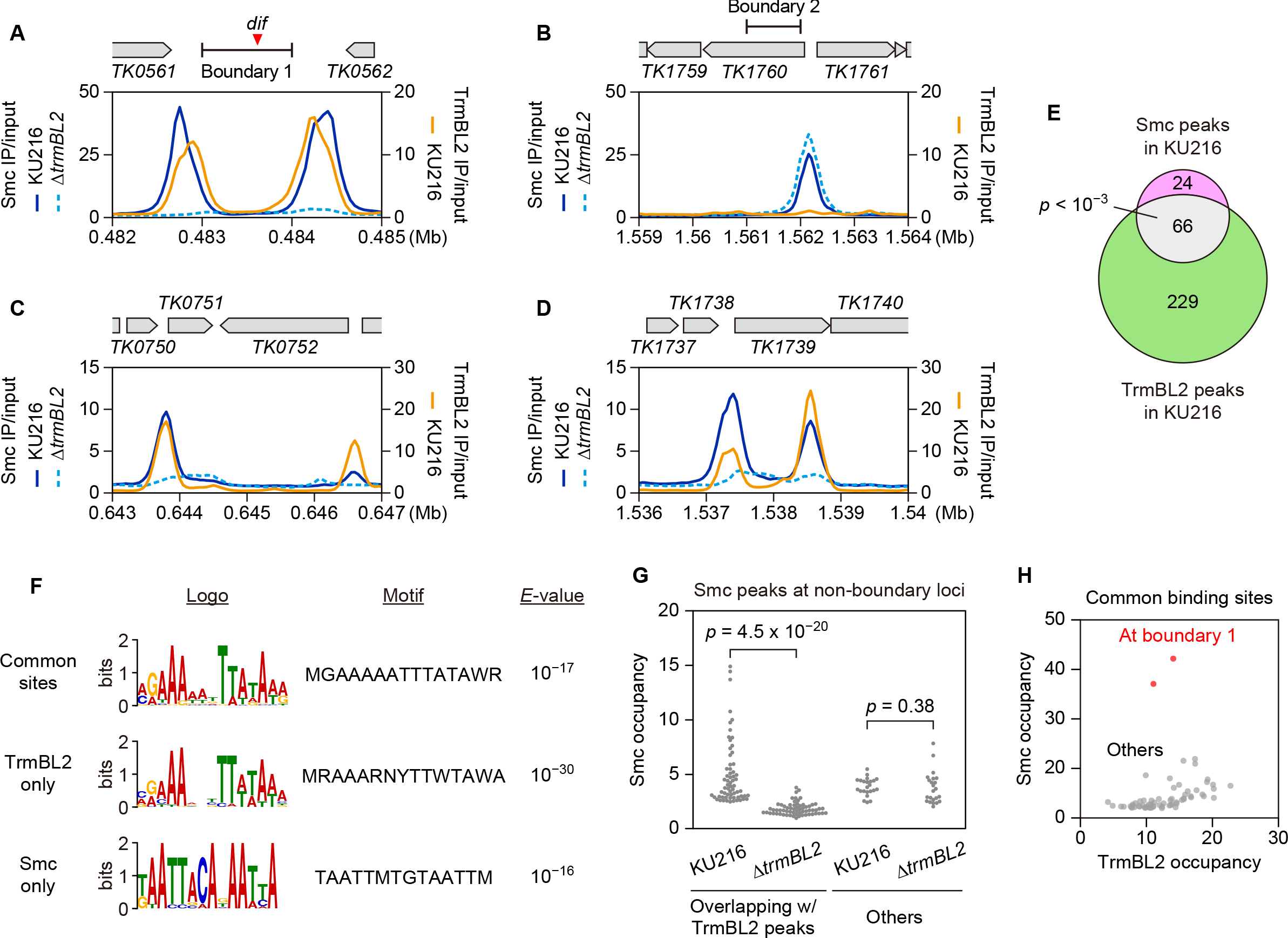
TrmBL2 is required to localize Smc-ScpAB to boundary 1 and non-boundary regions. (A-D) ChIP-seq tracks of Smc (in KU216 and Δ*trmBL2*) and TrmBL2 (in KU216) are shown for boundary 1 (A), boundary 2 (B), and non-boundary loci (C and D) in *T. kodakarensis*. Enrichment of immunoprecipitated versus input DNA (IP/input) is shown at 50-bp resolution. The *dif* position is indicated by a red triangle. The positions of boundaries 1 and 2 were determined at 1-kb resolution. Genes and their orientations are indicated by gray pentagons. (E) Venn diagram showing the overlap of Smc and TrmBL2 peaks detected by ChIP-seq analysis of KU216. Statistical significance of the overlap was determined by permutation test. (F) MEME-ChIP^97^ was performed to search for DNA motifs enriched in the three peak groups in (E). Statistical significance of the enrichment was evaluated using *E-*values. Only the most significant motif is shown for each group. (G) Impact of *trmBL2* deletion on Smc binding. Smc occupancy was calculated for non-boundary loci forming Smc peaks in KU216. Occupancy was defined as the ChIP-seq IP/input ratio of a 200- bp region centered at the peak summit. Statistical significance of the difference was determined by Wilcoxon rank sum test. (H) Occupancies of TrmBL2 and Smc were plotted for their common binding sites shown in (E).

At boundary 1, the peak summits of TrmBL2 were reproducibly located ∼100 bp inward from the cognate peak summits of Smc (Figures 6A and S6C). This distribution could be explained if TrmBL2, akin to CTCF, serves as an asymmetric barrier for loop-extruding Smc-ScpAB. ^58,59^

However, this scenario does not fit well with the crystal structure of DNA-bound *Pyrococcus furiosus* TrmBL2, in which two dimers of TrmBL2 interact with the DNA in a symmetric manner. ^60^ Another possibility is that each of the two closely located TrmBL2-binding sites at boundary 1 stalls Smc- ScpAB with high efficiency, possibly ∼100%. In this case, as loading of Smc-ScpAB onto the intervening region should be a relatively rare event, the left and right binding sites for TrmBL2 will encounter Smc-ScpAB mostly from the left and right, respectively. This directionally biased encounter and subsequent stalling will generate the observed ChIP-seq profile of Smc.

In KU216, a high ChIP-seq peak of Smc was also found near boundary 2 (Figures 6B and S6A). In contrast to boundary 1, however, this region was not enriched for TrmBL2 (Figures 6B and S6B). Moreover, deletion of *trmBL2* did not reduce the Smc binding to boundary 2. These results are consistent with a TrmBL2-independent formation of boundary 2 (Figures 5D and E).

Our ChIP-seq identified a total of 90 Smc peaks in KU216, among which 66 overlapped with TrmBL2 peaks in a statistically significant manner (Figures 6C-E). Common and TrmBL2-specific binding sites were enriched for very similar AT-rich motifs, whereas Smc-specific binding sites were characterized by a dissimilar AT-rich motif (Figure 6F). As observed for boundary 1, many non- boundary regions that were co-occupied by Smc and TrmBL2 in KU216 showed decreased binding of Smc in Δ*trmBL2* cells (Figure 6G). This was not due to global depletion of Smc from the chromosome, because the loss of TrmBL2 did not reduce Smc occupancy at Smc-specific binding sites (Figures 6B and G). Taken together, TrmBL2 stalls Smc-ScpAB at many non-boundary regions as well as boundary 1.

To explore the potential difference between boundary 1 and other stalling sites, we plotted the occupancies of TrmBL2 and Smc at their common binding sites (Figure 6H). The two values showed a good correlation on average. However, Smc was disproportionally enriched at a number of loci despite their modest enrichment for TrmBL2, and this disproportionality was most evident at boundary 1. The variability is also exemplified by the ChIP-seq profiles in Figures 6C and D, which display different heights of Smc peaks relative to the cognate TrmBL2 peaks. These results suggest that TrmBL2 blocks Smc-ScpAB very effectively at boundary 1 (possibly with ∼100% efficiency as described above) and much less, but still significantly, at non-boundary stalling sites. Importantly, this difference cannot simply be explained by the level of TrmBL2 occupancy.

### DNA sequences correlate with the efficiency of TrmBL2-dependent stalling of Smc-ScpAB

Why is the stalling efficiency of Smc-ScpAB variable between loci? We noticed that the intergenic region encompassing boundary 1 exhibited unusually high AT content (Figure 7A). The percentage was especially high around the peaks of Smc and TrmBL2, reaching a maximum of over 75% at 200- bp scale, while the overall AT content of the *T. kodakarensis* genome is 48%. AT content was also high at other loci co-occupied by TrmBL2 and Smc (Figures S7A and B). These observations led us to hypothesize that AT-rich sequences affect how efficiently TrmBL2 blocks loop-extruding Smc- ScpAB.

**Figure 7.**
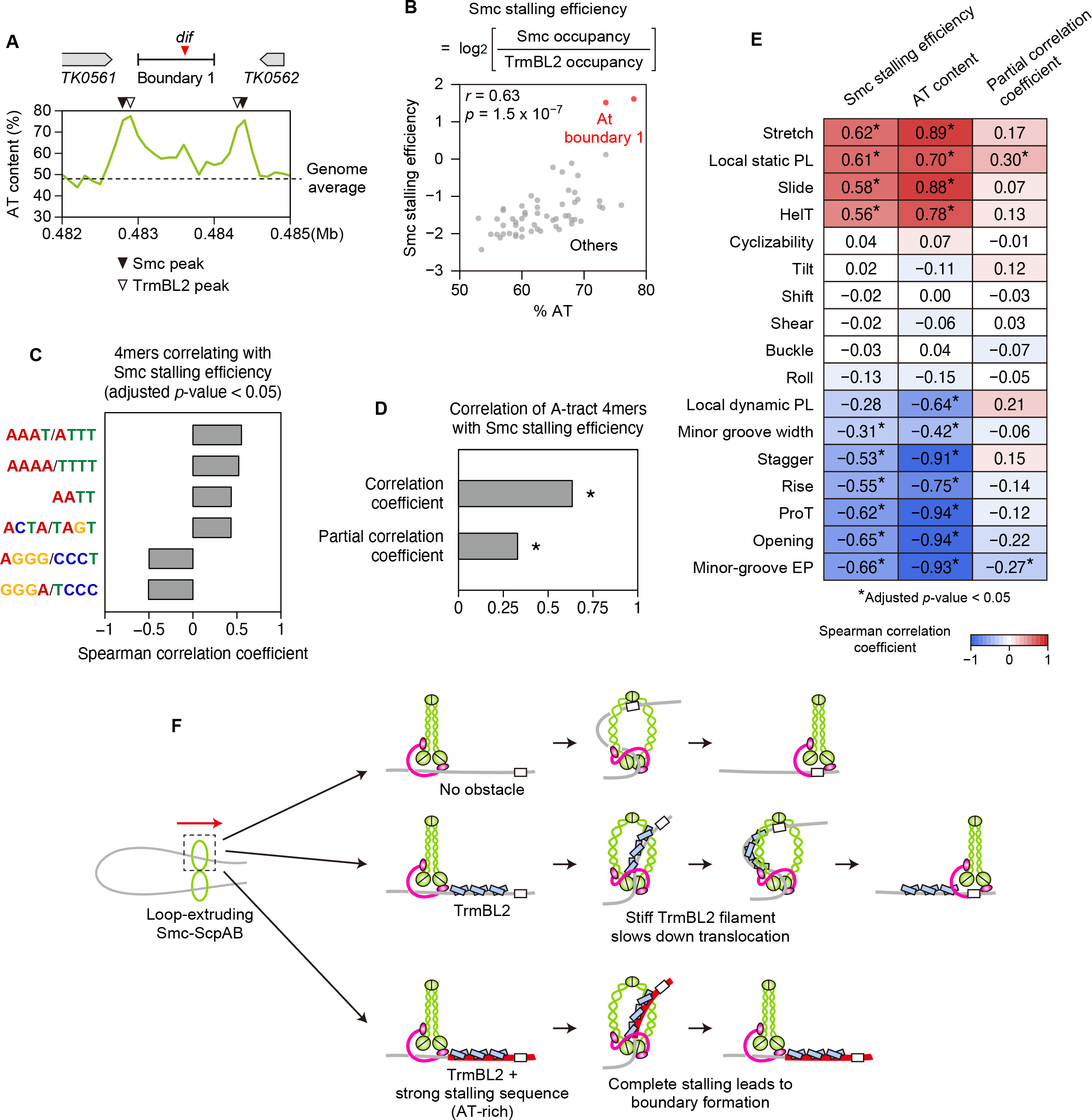
DNA sequence features correlating with TrmBL2-dependent stalling of Smc-ScpAB (A) AT-content track of boundary 1. AT content was calculated with sliding window size of 200 bp and step size of 100 bp. The peak positions of Smc and TrmBL2 are indicated by black and white triangles, respectively. (B) AT content and Smc stalling efficiency were plotted for 57 stalling sites. The Spearman rank correlation coefficient (*r*) and corresponding *p*-value are also shown. The two stalling sites associated with boundary 1 are highlighted in red. See STAR Methods for more detail. (C) Spearman correlation coefficients between DNA tetranucleotide frequencies and Smc stalling efficiency were calculated for the 57 stalling sites. Only the sequences with high statistical significance (adjusted *p-*value < 0.05) are shown. (D) The Spearman correlation coefficient between the frequency of the A-tract tetranucleotides and Smc stalling efficiency was calculated for the 57 stalling sites. The partial Spearman correlation coefficient was also calculated to examine their relationship without the contribution of AT content. An asterisk indicates a *p-*value lower than 0.05. (E) Spearman correlation coefficients of Smc stalling efficiency and AT content with DNA structural properties were calculated for the 57 stalling sites. Partial Spearman correlation coefficients between Smc stalling efficiency and DNA structural properties were calculated by removing the contribution of AT content. Asterisks indicate high statistical significance (adjusted *p-*value < 0.05). (F) Model for the TrmBL2-mediated regulation of Smc-ScpAB dynamics in *T. kodakarensis*. The diagram assumes that the complex functions as a dimer as suggested for bacterial Smc-ScpAB. ^23^ ScpB probably forms a complex with Smc (green) and ScpA (magenta) but is not included in the diagram for simplicity. See Discussion for more detail.

To explore this possibility, we defined a simple metric named Smc stalling efficiency. When a given TrmBL2-binding site overlapped with an Smc-binding site (each defined as a 200-bp region centered at the peak summit), their occupancy ratio log2(Smc/TrmBL2) was defined as the Smc stalling efficiency of the TrmBL2-binding site. If TrmBL2 stalls Smc-ScpAB more efficiently at a given locus than another, the value of Smc stalling efficiency is expected to be higher at the former. We calculated Smc stalling efficiency for the 57 TrmBL2-binding sites that overlapped with TrmBL2- dependent Smc-binding sites (see STAR Methods for more detail). These 57 loci can be seen as the most probable stalling sites. Comparison of Smc stalling efficiency with the AT content of the TrmBL2-bound 200-bp region revealed that the two metrics were well correlated with each other (Figure 7B). In contrast, AT content showed a weaker correlation with Smc occupancy and no significant correlation with TrmBL2 occupancy (Figures S7C and D). Thus, AT content is strongly linked to the stalling efficiency of Smc-ScpAB but not to the DNA binding affinities of Smc-ScpAB and TrmBL2 at stalling sites.

We next investigated whether Smc stalling efficiency is associated with more complex sequence features. We counted occurrence of each of all possible DNA tetranucleotides in the TrmBL2-bound 200-bp regions used for the calculation of Smc stalling efficiency. The Spearman correlation coefficient was calculated between the tetranucleotide frequency and Smc stalling efficiency. This analysis identified all types of so-called “A-tract” tetranucleotides (AAAA/TTTT, AAAT/ATTT, and AATT/AATT) as the most positively correlated sequences (Figure 7C). A-tracts are four or more consecutive A:T base pairs without a TpA step. ^61–63^ The total number of the A-tract tetranucleotides showed a significant correlation with Smc stalling efficiency even if the contribution of AT content was removed by partial correlation analysis (Figure 7D). This indicates a potential role of A-tracts in helping TrmBL2 to stall Smc-ScpAB.

A-tracts display unique structural features in multiple ways. For example, long poly(A) sequences (often 10-20 bp or even longer) disfavor nucleosome wrapping in eukaryotes, while A-tracts induce DNA curvature when in phase with the helical pitch of DNA. ^63–65^ A-tracts also form narrow and negatively charged minor grooves. ^66^ The overrepresentation of A-tracts in efficient stalling sites raises the possibility that TrmBL2-dependent stalling of Smc-ScpAB is affected by structural properties of the underlying sequence. To test this, we quantified various features of stalling sites solely by using their sequence information. We first focused on persistence length (PL), within which a polymer segment can be seen as a straight rod. Although the PL of DNA is often regarded as its stiffness to bending deformation, the metric can actually be decomposed into dynamic and static PLs, which reflect *bona fide* stiffness and intrinsic shape of the DNA chain, respectively. ^67,68^ These two types of PLs have recently been estimated for all possible DNA tetranucleotides using all-atom molecular dynamics (MD) simulation data. ^69^ We used these values to calculate the mean dynamic and static PLs of all tetranucleotide steps in each stalling site, seeing the values as the local PLs of the site. We also predicted the DNA “cyclizability” of each stalling site using DNAcycP. ^70^ This deep- learning-based tool relies on the experimentally measured efficiency of DNA looping mediated by complementary single-stranded overhangs of linear DNA. ^71^ Lastly, we used the recently developed software Deep DNAshape^72^ to predict groove features (width and electrostatic potential [EP] of the minor groove), intra-base-pair features (Shear, Stretch, Stagger, Buckle, Propeller Twist [ProT], and Opening), and inter-base-pair features (Shift, Slide, Rise, Tilt, Roll, and Helical Twist [HelT]) averaged over the bases of each sequence. Among these 17 structural features, 11 showed significant correlation with both Smc stalling efficiency and AT content (Figure 7E). When partial correlation analysis was performed to remove the contribution of AT content, 9 of the 11 features displayed no significant correlation with Smc stalling efficiency. These nine features may thus be simply associated with AT content rather than with efficient stalling of Smc-ScpAB. Meanwhile, the other two features (local static PL and minor groove EP) exhibited significant positive and negative correlations with Smc stalling efficiency respectively even after the contribution of AT content was removed. Thus, these two features are indeed associated with Smc stalling efficiency and may be causally related to it.

Current models posit that DNA bending is prerequisite for loop-extruding SMC complexes to reel in DNA for translocation. ^7,73,74^ Given this, it is somewhat surprising that local dynamic PL did not significantly contribute to Smc stalling efficiency. Experimental evidence suggests that SMC complexes capture a bent DNA segment of ∼200 bp to carry out a loop extrusion cycle. ^75–78^ The effect of DNA stiffness may need to be evaluated at this length scale. This prompted us to estimate the dynamic PLs of entire 200-bp segments (here referred to as gross dynamic PL) of stalling sites by performing coarse-grained MD simulations. ^79^ To save the computational cost for the modeling, we selected and analyzed the 5 most efficient and 5 most inefficient stalling sites among the 57 stalling sites. The directional correlation decay of the modeled DNA polymer was used to calculate gross dynamic PL. The obtained value actually showed a high positive correlation with Smc stalling efficiency (*r* = 0.85)(Figure S7E). However, the gross dynamic PL varied only from 64.8 to 68.1 nm among the ten sequences, whereas the local static PL varied in a much larger range of 88.0 to 211 nm among the 57 stalling sites (Figure S7F). At this stage, it remains unclear whether the small differences in the gross dynamic PLs contribute to the variability in Smc stalling efficiency. It is possible that our MD simulation analysis has underestimated the gross dynamic PL.

## DISCUSSION

Our genetic analysis has demonstrated that Smc, ScpA, and ScpB are all required for the domain formation in *T. kodakarensis* (Figure 2). This is consistent with the dogma that canonical SMC proteins form holocomplexes with Kleisin and its partner Kite/Hawk subunits. ^1,4,5^ However, Jeon et al. failed to detect a physical interaction between archaeal ScpA and ScpB *in vitro*. ^30^ In contrast to the case with bacteria, ^32,33^ archaeal ScpA and ScpB may require an additional protein for a detectable interaction. Indeed, our analyses using yeast two-hybrid assays and modeling by AlphaFold2^80^ suggest that archaeal ScpA and ScpB can form a complex when Smc triggers a conformational change of ScpA (Takemata et al., manuscript in preparation).

In *T. kodakarensis*, TrmBL2 is required to localize Smc-ScpAB to tens of loci, most strikingly at boundary 1 (Figure 6). We propose that TrmBL2, analogous to eukaryotic CTCF, ^6,10,59^ stalls loop- extruding Smc-ScpAB and that the stalling occurs most frequently at the *dif*-proximal TrmBL2- binding sites, resulting in the formation of boundary 1 (Figure 7F). Assuming high stalling efficiencies (possibly close to 100%) for the TrmBL2-enriched regions at boundary 1, this model can explain the distribution pattern of Smc observed at boundary 1 (Figures 6A and S6C). A potential alternative explanation for the colocalization of Smc and TrmBL2 is that TrmBL2 serves as a loader of Smc-ScpAB. A simulation study showed that, if a loop-extruding factor shapes a boundary at its loading site as postulated above, it must perform asymmetric loop extrusion combined with one- dimensional diffusion along DNA. ^81^ However, although the loop extrusion of Smc-ScpAB has not been directly observed *in vitro*, Hi-C studies suggested that the complex serves as a dimer that extrudes a DNA loop symmetrically. ^23^ Single-molecule experiments also demonstrated that the Kite- containing eukaryotic SMC complex SMC5/6 and the bacterial MukBEF-like complex Wadjet form dimeric supercomplexes to extrude DNA loops symmetrically. ^82,83^ Taken together, it is more plausible that *T. kodakarensis* Smc-ScpAB is also a symmetric loop extruder and is stalled by TrmBL2 for the boundary formation.

Our study highlights fundamental differences in the Smc-mediated genome organization between archaea and bacteria. First, the above model places *T. kodakarensis* Smc-ScpAB as a key player for a eukaryotic-like mechanism of domain formation, whereas bacterial Smc-ScpAB is not involved in sculpting domains. ^18,19^ Instead, bacterial Smc-ScpAB promotes sister chromosome resolution by being recruited to the replication origin and juxtaposing the two intra-chromosomal arms through loop extrusion. ^23,25–27^ The chromosomal loading of bacterial Smc-ScpAB is mediated by the DNA-binding CTPase ParB, which binds to *parS* centromere-like sequences scattered around the origin. ^25,26,84–86^ Notably, the *T. kodakarensis* homolog of ParB functions as an ADP-dependent serine kinase involved in cysteine biogenesis, and thus this enzyme does not appear to play a role in Smc-ScpAB loading. ^87^

It is interesting that boundary 1 is located near *dif*, a highly conserved *cis* element contributing to the genome integrity in many prokaryotes (Figure 3). *dif* and *dif-*like sequences are also found proximal to boundary structures in *T. acidophilum*, *H. volcanii*, and *H. salinarum*. All of these organisms possess both Smc proteins and TrmBL2-like proteins (data not shown). These findings suggest that the mechanism and function of the *dif-*associated boundary are conserved in multiple euryarchaeal lineages. However, the physiological consequences of *dif-*associated boundaries remain unclear due to the fact that we have not seen a clear phenotype associated with defective formation of boundary 1. *dif*-associated boundaries may play a role under specific conditions, for example when the Xer/*dif* system becomes more critical due to increased DNA damage.

According to current models, the loop extrusion cycle of SMC complexes requires a DNA bend to reel a DNA segment through the lumen of the SMC ring. ^7,73,74^ We propose that TrmBL2, a NAP that forms a filamentous structure along DNA, serves as an obstacle that blocks loop extrusion by slowing or prohibiting DNA bending (Figure 7F). A previous biophysical study demonstrated that TrmBL2 increases the persistence length of bound DNA in a concentration-dependent manner. ^56^ The persistence length can be increased even above 90 nm, approximately twice as large as the size of the SMC ring. In line with our model, a recent study observed that the loop extrusion of eukaryotic condensin is stalled by a linear array of Rap1, a telomeric protein that can induce local DNA stiffening.^88,89^

To our surprise, we found that TrmBL2-dependent stalling of Smc-ScpAB is associated with the AT richness of the underlying sequence (Figure 7B). Given the high growth temperature of *T. kodakarensis* (85°C), one could imagine that AT-rich sequences may interfere with loop extrusion by being denatured into single-stranded DNA (ssDNA). This form of DNA has been suggested to serve as a high-affinity substrate for Smc-ScpAB binding and thereby may trap the complex. ^90^ However, due to the fragility of ssDNA that threatens genome integrity, hyperthermophiles are more likely to suppress heat-induced denaturation of genomic DNA. For this suppression, all hyperthermophiles appear to leverage reverse gyrase, a type IA topoisomerase that generates positive DNA supercoils and is critical for the thermal adaptation of hyperthermophiles. ^91–93^ Hyperthermophilic archaea also protect their genomes from thermal denaturation by coating genomic DNA with NAPs more extensively than mesophilic archaea do. ^94^ We prefer the idea that physical properties of AT-rich sequences other than their proneness to denaturation facilitate TrmBL2-mediated stalling of Smc-ScpAB. This notion is supported by the positive correlation of efficient stalling with A-tracts, a specific type of AT-rich sequences that displays unique structural features (Figures 7C and D). A-tracts are characterized by their intrinsically straight conformation and enhanced minor-groove EP. ^63,69,95^ It is plausible that these two features of A-tracts help TrmBL2 to impede Smc-ScpAB translocation, which is consistent with their correlations with Smc stalling efficiency (Figure 7E). We also note that the DNA motifs found at TrmBL2- and Smc-binding sites contain A-tract-like sequences (Figure 6F). The structural properties of A-tracts may affect their interaction with TrmBL2 or Smc-ScpAB, thereby influencing the stalling competency. In this case, the effect of A-tracts on DNA binding of TrmBL2 or Smc-ScpAB will be qualitative rather than quantitative, given that AT richness is not markedly correlated with the binding level of TrmBL2 or Smc-ScpAB (Figures S7C and D). Minor-groove EP can indeed affect protein-DNA interaction via so-called shape readout mechanisms. ^66,95^

In summary, our study has not only uncovered a eukaryotic-like mechanism of chromosomal domain formation in archaea, but also shed light on the potential role of intrinsic DNA structure in defining higher-order genome organization. It will be interesting to see whether a similar interplay of SMC complexes, boundary proteins, and structural features of boundary sequences shapes eukaryotic 3D genomes.

## Limitations of the study

3C-seq data of *T. kodakarensis* could contain a significant fraction of inter-chromosomal contacts due to its polyploidy, but they cannot be distinguished from intra-chromosomal contacts. Our 3C-seq data also contain uncertainty resulting from the genome heterogeneity caused by the inversion. Currently, the physiological consequences of the chromosomal domains are unclear. It should be emphasized that the boundary formation mechanism proposed here is inapplicable to all types of domain boundaries in archaea, as exemplified by the TrmBL2-independent formation of boundary 2. Future studies will be required to dissect what structural features of AT-rich sequences impact TrmBL2- mediated stalling of Smc-ScpAB.

## ACKNOWLEDGMENTS

We thank Single-Cell Genome Information Analysis Core (SignAC) at WPI-ASHBi at Kyoto University for Illumina DNA sequencing. We thank Takeshi Yamagami and Suzuka Kajikawa for their technical assistance to purify the Smc protein, Itaru Hamachi and Tomonori Tamura for mass spectrometry analysis, Hugo Maruyama for his kind gifts of the *T. kodakarensis* KCP1 strain and the TrmBL2 expression plasmid, and Stephen Bell for helpful comments on the manuscript. The *T. acidophilum* strain DSM 1728 (JCM 9062) was provided by Japan Collection of Microorganisms at RIKEN BRC, which is participating in the National BioResource Project of the MEXT, Japan. The supercomputer of Academic Center for Computing and Media Studies (Kyoto University) was used for the MD simulations. K.Y. is supported by JST, the establishment of university fellowships towards the creation of science technology innovation, Grant Number JPMJFS2123. N.T. is funded by JST PRESTO (JPMJPR20K7), JST FOREST (JPMJFR224V), JSPS Grants-in-Aid for Young Scientists Start-up (JP21K20636), JSPS Grants-in-Aid for Transformative Research Areas (JP23H04281), Takeda Science Foundation, and the Uehara Memorial Foundation. S.T. is funded by JSPS Grants-in-Aid for Transformative Research Areas (JP20H05934). Y.I. is funded by JSPS Grant-in-Aid for Challenging Research (Exploratory)(JP19K22289).

## AUTHOR CONTRIBUTIONS

K.Y. and K.M. performed 3C-seq and strain construction *for T. kodakarensis*. K.Y. also performed growth measurement, RNA-seq, and ChIP-seq. N.T. performed 3C-seq on *T. acidophilum* and DNA affinity purification. K.Y. and S.I. performed protein purification to raise antibodies for ChIP-seq.

N.T. performed data analysis on DNA sequencing data. M.Y. and S.T. performed MD simulations.

N.T. and H.A. supervised the project. N.T., S.T., and Y.I. acquired funding. K.Y. and N.T. wrote the manuscript. All co-authors critically read and edited the manuscript.

## DECLARATION OF INTERESTS

The authors declare no competing interests.

## STAR★METHODS

### KEY RESOURCES TABLE

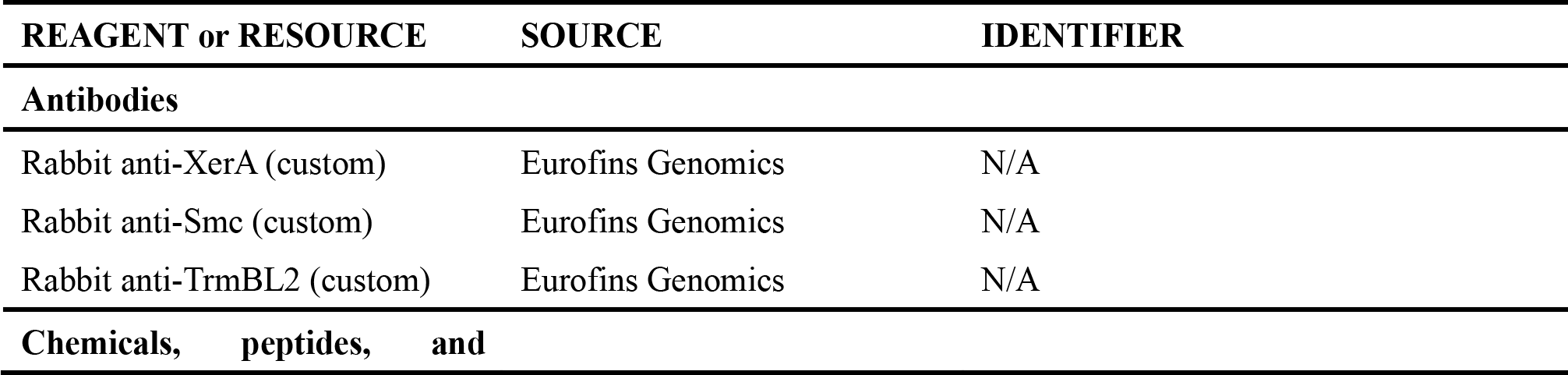
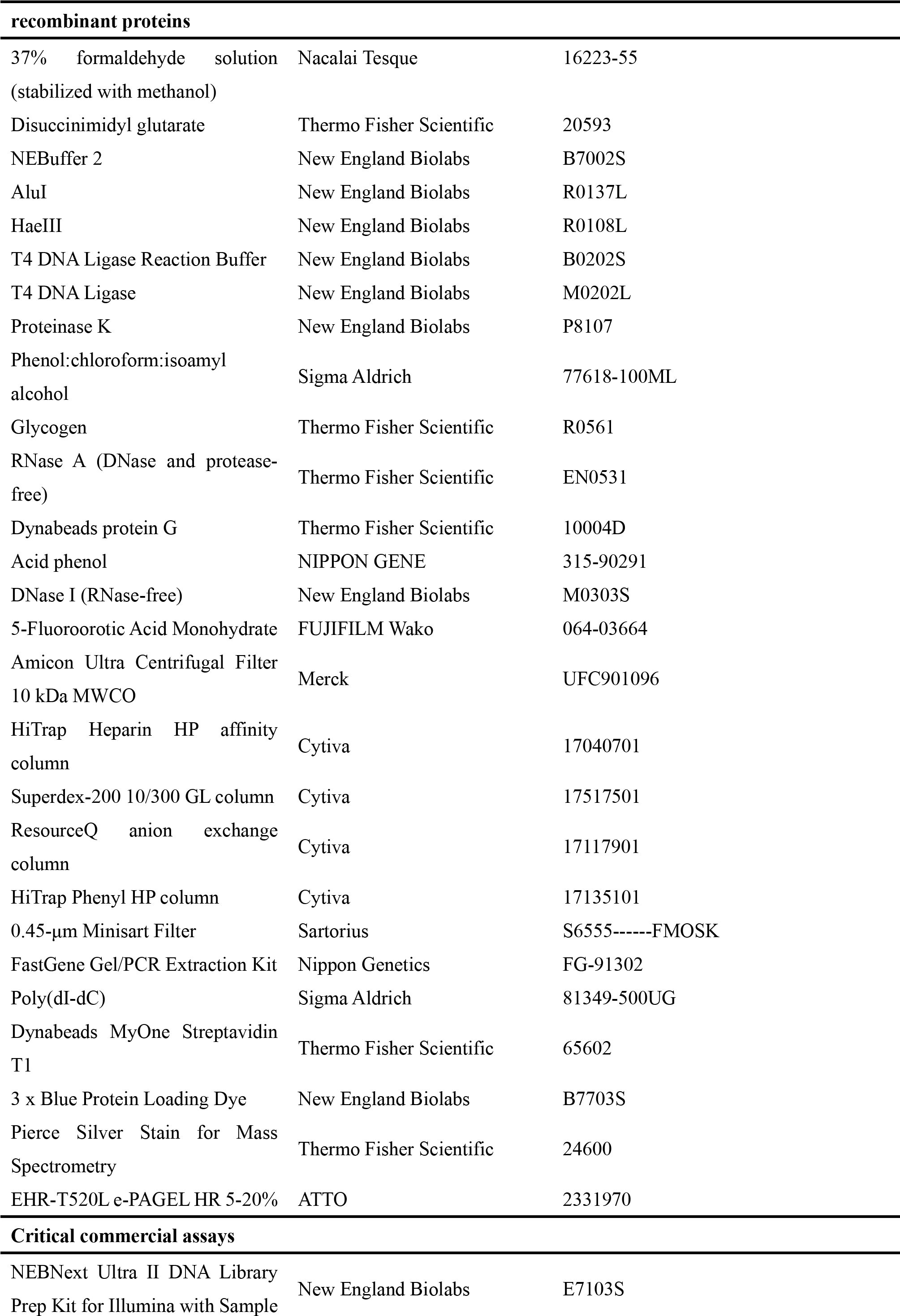
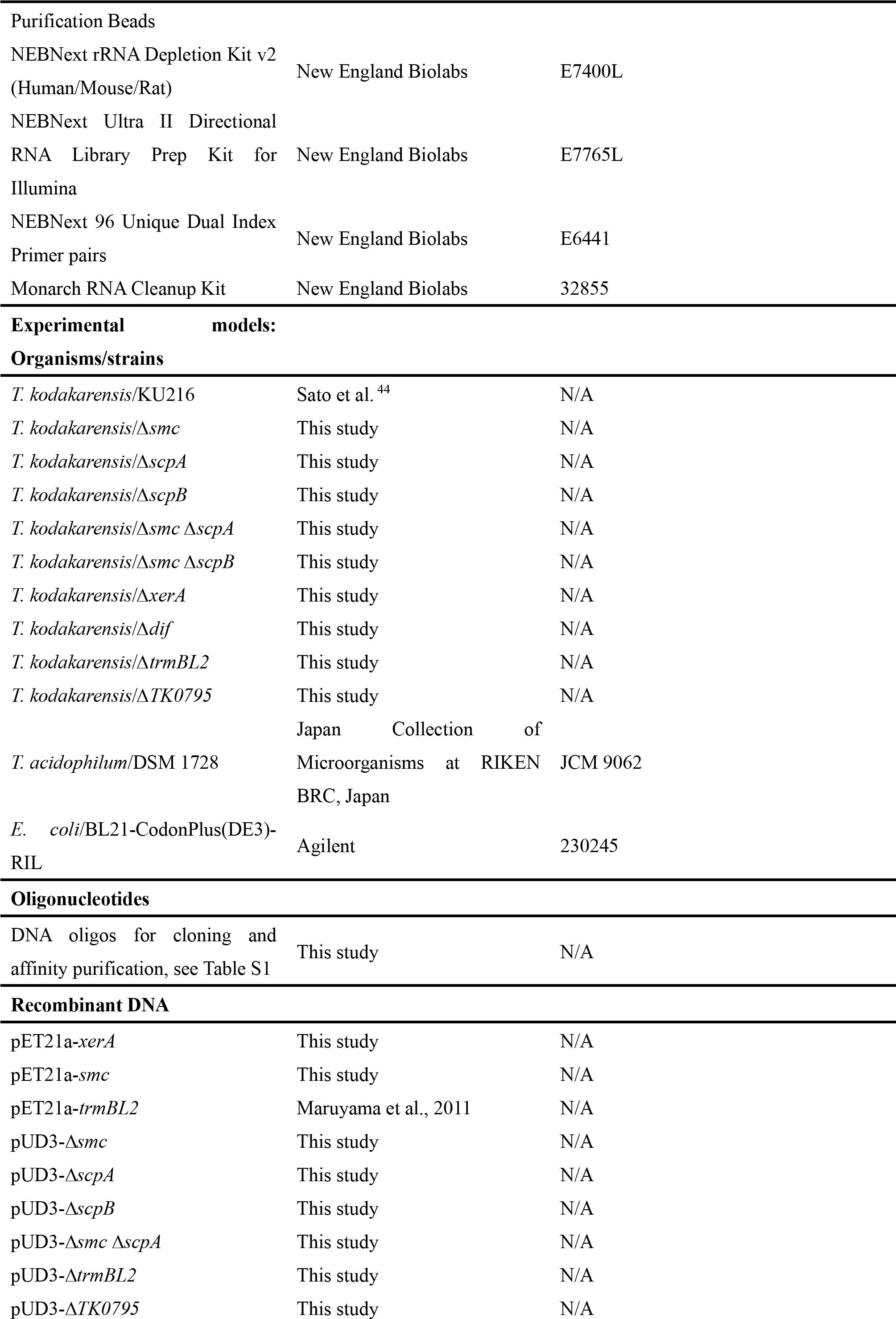
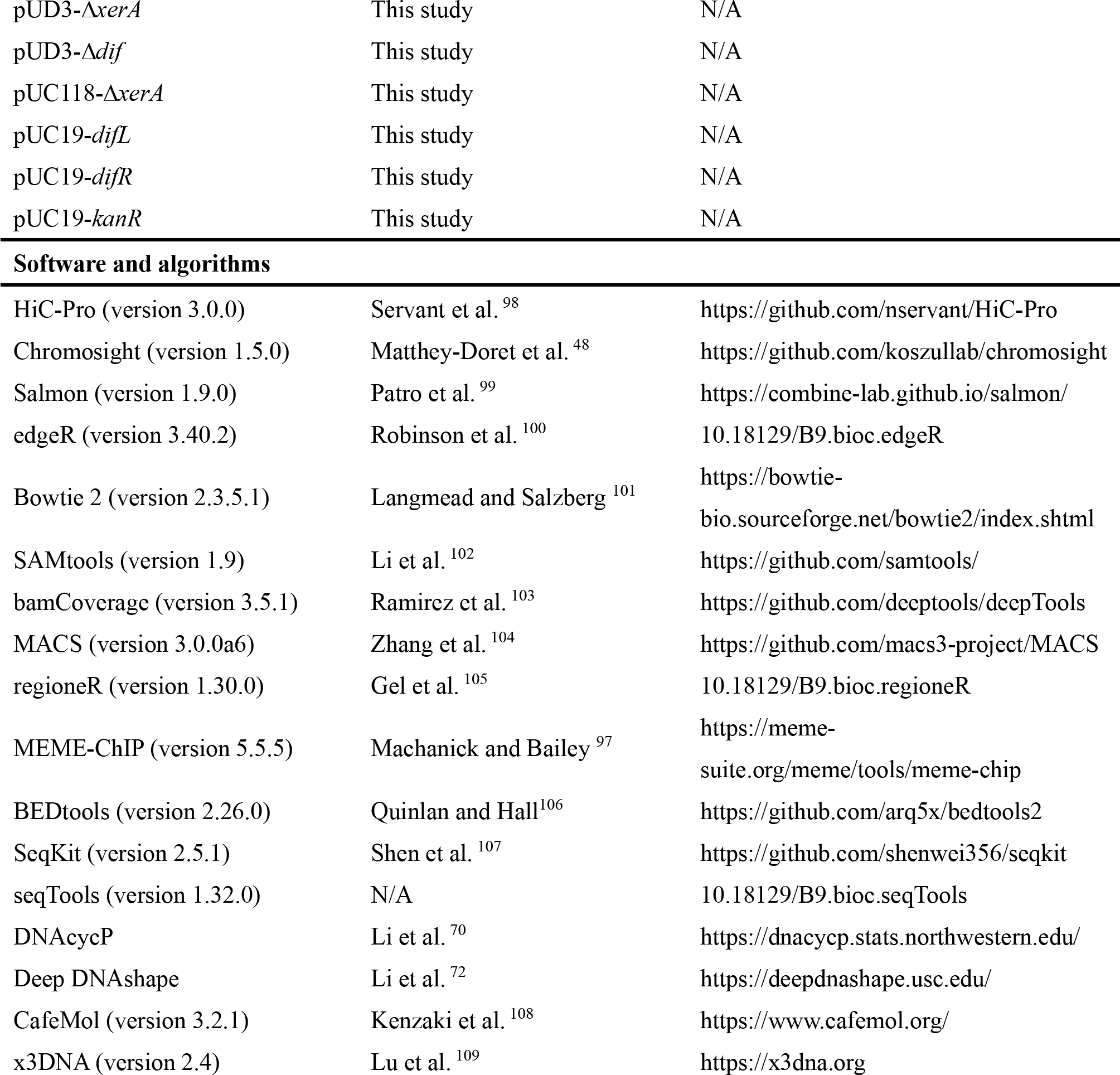

### RESOURCE AVAILABILITY

#### Lead contact

Further information and requests for resources and reagents should be directed to and will be fulfilled by the lead contact, Naomichi Takemata.

#### Materials availability

*T. kodakarensis* strains and antibodies generated for this study are available upon request to the lead contact.

#### Data and code availability

This paper does not report original code. Any additional information required to reanalyze the data reported in this paper is available from the lead contact upon request.

### EXPERIMENTAL MODEL AND SUBJECT DETAILS

#### T. kodakarensis

The uracil-auxotrophic strain KU216 (*ΔpyrF*) ^44^ was used as a parental strain for strain construction. For all experiments, cells were grown anaerobically at 85°C. Unless otherwise stated, cells were cultivated as follows. Cells were inoculated into the ASW-YT-m1-S^0^ rich medium^110^ and pre-cultured overnight. The pre-culture was inoculated into fresh ASW-YT-m1-S^0^ rich medium to an OD660 of 0.015. The culture was grown until mid-to-late log phase (OD660: 0.2-0.25) and used for experiments.

ASW-YT-m1-S^0^ was used with the redox indicator resazurin added to a concentration of 0.5 mg/L.

#### T. acidophilum

The *T. acidophilum* wild-type strain DSM 1728 was grown as described previously^53^ to an OD600 of ∼0.2.

#### E. coli

The BL21-CodonPlus(DE3)-RIL strain (Agilent) was used for protein expression. Unless otherwise stated, cells harboring an expression plasmid were grown in LB medium containing 100 μg/mL of ampicillin and 30 μg/mL of chloramphenicol.

### METHOD DETAILS

#### Construction of *T. kodakarensis* strains

Except *xerA* (*TK0777*), all genes were deleted using a pop-in/pop-out approach targeting the entire (or almost entire) coding region. ^44^ For *xerA* deletion, its coding sequence was replaced with a marker gene via double crossover. ^111^ For *dif* deletion, the entire 28-bp sequence of *dif* (TTTTGATATAATGTACCTTATATGACAA) was deleted using the pop-in/pop-out method.

#### Deletion constructs

The primers used for the construction are listed in Table S1. For deletions of *smc* (*TK1017*), *scpA* (*TK1018*), and *scpB* (*TK1962*), each target sequence was PCR amplified together with upstream and downstream ∼1-kb regions and cloned into the pUD3 plasmid, which contains the *pyrF* marker. ^112^ pUD3 was digested at the PstI and SalI sites for the cloning. The target sequence was then removed by inverse PCR to generate the deletion plasmids pUD3-Δ*smc*, pUD3-Δ*scpA*, and pUD3-Δ*scpB*. For double deletion of *smc* and *scpA*, constituting a part of an operon, pUD3-Δ*smc* was amplified by inverse PCR to remove the downstream homology arm. The obtained PCR fragment was fused with a downstream ∼1-kb region of *scpA* using In-Fusion HD Cloning Plus reagents (Takara Clontech) to generate pUD3-Δ*smcΔscpA*. For *dif* deletion, upstream and downstream ∼1-kb regions of the *dif* sequence were PCR amplified separately and cloned together into pUD3 (digested at BamHI and HindIII sites) using In-Fusion HD Cloning Plus reagents. The obtained plasmid was named pUD3-Δ*dif*. For *TK0795* deletion, upstream and downstream ∼1-kb regions of the target sequence were PCR amplified separately and cloned together into the EcoRI site of pUD3 using In-Fusion HD Cloning Plus reagents. The obtained plasmid was named pUD3-Δ*TK0795*. For *trmBL2* (*TK0471*) deletion, upstream and downstream ∼1-kb regions of the target sequence were PCR amplified as a single fragment using genomic DNA of the KCP1 strain^55^ (Δ*pyrF* Δ*trpE* Δ*trmBL2*) as a template. The KCP1 strain was kindly provided by Hugo Maruyama. The fragment was cloned into the EcoRI site of pUD3 to generate the deletion plasmid pUD3-Δ*trmBL2*. For *xerA* deletion, a PCR fragment containing the *pyrF* marker and upstream and downstream ∼1-kb regions of the target sequence were PCR amplified separately. The three fragments were cloned together into the EcoRI site of pUC118 using In-Fusion HD Cloning Plus reagents. The obtained deletion plasmid was named pUC118-Δ*xerA*.

#### Transformation

For gene disruption via pop-in/pop-out recombination, transformation with deletion constructs described above was performed as described previously^110^. The synthetic medium ASW-AA-m1-S^0^ was made according to Su et al.. ^113^ For selection of Δ*trmBL2* cells, cells were spread onto solid ASW- AA-m1-S^0^ medium containing 7.5 g/L 5-fluoroorotic acid, 44 mM NaOH, and 5 mg/L uracil. The Δ*smc* Δ*scpA* strain was constructed by transforming KU216 with pUD3-Δ*smc*Δ*scpA*. The Δ*smc* Δ*scpB* strain was constructed by disrupting *scpB* in the Δ*smc* mutant.

Double-crossover deletion of *xerA* was conducted as follows. KU216 cells from 20 mL of an overnight culture were pelleted and resuspended in 200 µL of 0.8 x ASW-m1, which is composed of 0.8 x artificial seawater (ASW) ^114^ supplemented with 20 μM KI, 20 μM H3BO3, and 10 μM NiCl2. The cell suspension was mixed with 3 μg of the *xerA* deletion construct and incubated on ice for 5 min. The cells were heat shocked for 45 sec at 85°C and cooled on ice for 5 min. This heating-chilling cycle was repeated five times. The cell suspension was inoculated into 20 mL of the uracil-free ASW- AA-m1-S^0^ medium and grown for three days. 400 µL of the culture were inoculated into fresh ASW- AA-m1-S^0^ medium and further grown for three days. For single colony isolation, 50 µL of the culture were spread onto ASW-YT-m1 medium solidified with 10 g/L Gelrite and grown for one day. Deletion of *xerA* was confirmed by colony PCR and DNA sequencing.

### Growth measurement

For each replicate, a pre-culture was inoculated into three vials containing fresh ASW-YT-m1-S^0^. These three cultures were grown at 85°C and used separately for OD660 measurement.

### 3C-seq

#### Cell Fixation

*T. kodakarensis* cells were crosslinked by dispensing 8 mL of a hot culture into each of eight 50-mL tubes containing 32 mL of fixative solution 1 (4.3 ml of 37% formaldehyde, 6.4 mL of 4 x ASW-m1, and 21.3 ml of MilliQ water). The mixtures were incubated for 30 min at 25°C with agitation at 75 rpm. Crosslinking was quenched by adding 10 mL of 2.5 M glycine to each tube and incubating the mixtures for 10 min at room temperature. The cells were collected by centrifugation (8,000 x g, 15 min, 25°C). Supernatant was removed with ∼2 ml per tube left for resuspension. The cell suspensions were combined and dispensed into four 5-mL tube. The cells were centrifuged (10,000 x g, 5 min, 25 °C), and two each of the pellets were resuspended in 3 mL of fixative solution 2 (30 µL of 300 mM DSG [Thermo Fisher Scientific] dissolved in DMSO, 600 µL of 4 x ASW-m1, and 2370 µL of MilliQ water) for additional crosslinking. The two cell suspensions were incubated for 40 min at 25°C with agitation at 75 rpm. Crosslinking was quenched by adding 750 µL of 2.5 M glycine and incubating for 5 min at room temperature. The cells were collected by centrifugation (10,000 x g, 5 min, 4°C). The two pellets were resuspended together in 1 mL of 1 x PBS. The cells were centrifuged (10,000 x g, 5 min, 4°C), resuspended in 1 mL of 1 x PBS, and centrifuged again (10,000 x g, 5 min, 4°C). The pellet was stored at –80°C until use.

For crosslinking of *T. acidophilum*, 50 mL of cell culture were pelleted (8,000 x g, 5 min, room temperature) and resuspended in a mixture of 3.35 mL 1 x PBS and 0.65 mL 37% formaldehyde (final 6%). The reaction was incubated for 30 min at 25°C with agitation at 100 rpm. Crosslinking was quenched by adding 680 µL of 2.5 M glycine and incubating the mixture for 10 min at room temperature. The cells were pelleted by centrifugation (10,000 x g, 3 min, 4°C). The centrifugation was repeated with the opposite side of the tube outward. The cells were washed by resuspending the pellet in 1 mL of 1 x PBS and centrifuging the suspension (10,000 x g, 2 min, 4°C). This step was repeated once. The pellet was stored at –80°C until use.

#### Restriction digestion

A frozen pellet of *T. kodakarensis* was resuspended in wash buffer (10 mM Tris-HCl [pH 8.0], 50 mM NaCl, 10 mM MgCl2) to an OD660 of 3. 1.2 mL of the cell suspension was used for the experiment. A frozen pellet of *T. acidophilum* was resuspended in 1 mL of wash buffer, and the whole cell suspension was used for the experiment. The cells were centrifuged (10,000 x g, 10 min, 4°C) and resuspended in 50 μL of 1 x NEBuffer 2 (New England Biolab). 50 μL of the suspension was transferred to an 1.5-mL tube, mixed with 5.55 μL of 10% SDS, and incubated for 15 min at 65°C with agitation at 600 rpm. The suspension was immediately cooled on ice for 90 sec and placed at room temperature until use. 12.5 µL of the lysate were stored for purification of undigested control DNA. For DNA digestion, 37.5 µL of the lysate was mixed with 232.5 µL of premixed buffer composed of 26.3 µL of 10 × NEBuffer 2, 60 µL of 10% v/v Triton X-100, and 146.2 μL of MilliQ water. This was further mixed with 15 μL each of 10 U/μL HaeIII (New England Biolabs) and 10 U/μL AluⅠ (New England Biolabs) and incubated for 3 h at 37°C with agitation at 600 rpm. The tube was also inverted every 30 min for mixing. After the incubation was completed, the reaction was mixed with 33.3 μL of 10% SDS and incubated for 10 min at room temperature to inactivate the restriction enzymes.

#### Proximity ligation

A previous study demonstrated that, for prokaryotic Hi-C, using only the insoluble material for proximity ligation significantly improves the data quality. ^38^ According to this strategy, the insoluble fraction was collected by centrifugation (16,000 x g, 10 min, 25°C), resuspended in 500 μL of wash buffer, and centrifuged again (16,000 x g, 10 min, 25°C). The pellet was resuspended in 1,185 μL of 1.01 x T4 DNA Ligase Reaction Buffer (New England Biolabs). 395 µL of the sample were mixed with 5 µL of MilliQ water and stored for purification of un-ligated control DNA. For ligation, 790 µL of the sample were mixed with 10 μL of 400 U/μL T4 DNA Ligase (New England Biolabs) and incubated for 3 h at 16°C with agitation at 600 rpm. The reaction was also mixed every 30 min by inverting the tube.

#### Reverse crosslinking

The insoluble fraction was collected by centrifugation (16,000 x g, 10 min, 25°C) and resuspended in 115 μL (for *T. kodakarensis*) or 230 µL (for *T. acidophilum*) of decrosslinking buffer (100 µL of TE buffer [pH 8], 10 µL of 10% SDS, and 5 µL of 0.5 M EDTA [pH 8]). The suspension was mixed with 3 μL (for *T. kodakarensis*) or 6 µL (for *T. acidophilum*) of 800 U/mL proteinase K (New England Biolabs). For *T. kodakarensis*, decrosslinking was performed by incubating the sample overnight at 37°C with agitation at 600 rpm. After that, 3 μL of 800 U/mL proteinase K were additionally added, and the sample was incubated for 4 h at 37°C with agitation at 600 rpm. For *T. acidophilum*, decrosslinking was performed by incubating the sample for 6 h at 65°C with agitation at 600 rpm, which was followed by overnight incubation at 30°C with agitation at 600 rpm. After that, 6 µL of 800 U/mL proteinase K were additionally added, and the sample was incubated for 4 h at 30°C with agitation at 600 rpm.

#### Sonication and purification of DNA

90 μL of the sample were used for DNA sonication, while the remainder of the sample was stored for purification of unsonicated control DNA. The 90-µL aliquot was transferred to a microTUBE AFA Fiber Pre-Slit Snap-Cap 6 x 16 mm (Covaris) and sonicated for 200 sec at 7°C using M220 Focused- ultrasonicator (Covaris) with the peak power set to 50, the duty factor set to 20, and the cycles/burst set to 200. DNA was then extracted twice with phenol:chloroform:isoamyl alcohol (Sigma Aldrich) and ethanol-precipitated together with 2 μL of 20 mg/mL glycogen (Thermo Fisher Scientific). The DNA was dissolved in 30 μL of 10 mM Tris-HCl (pH 8.0) containing 0.05 mg/mL RNase A (Thermo Scientific).

#### Library construction

The sheared DNA was used for library construction with NEBNext Ultra II DNA Library Prep with Sample Purification Beads (New England Biolabs). Reaction was performed according to the manufacturer’s instructions with size selection for a 300-400 bp insert.

### RNA-seq

#### RNA extraction

10 mL of a cell culture were dispensed into a 50-mL tube containing 20 mL of pre-chilled 0.8 x ASW- m1 and further chilled in iced water for 5 min. The cells were centrifuged (10,000 x g, 4°C, 10 min), resuspended in 1 mL of pre-chilled 0.8 x ASW-m1, and centrifuged again (10,000 x g, 4°C, 5 min). The pellet was resuspended in 750 µL of lysis buffer (75 µL of 10% SDS, 3 M sodium acetate [pH 5.2], and 650 µL of RNase-free water). The RNA was extracted twice with acid phenol (NIPPON GENE) and isopropanol-precipitated. The pellet was dissolved in 51 µL of RNase-free water. The solution was mixed with 6 µL of 10 x DNase I Reaction Buffer (New England Biolabs) and 3 µL of 2,000 U/mL DNase I (New England Biolabs). After 30 min of incubation at 37°C, the RNA was extracted with acid phenol and ethanol-precipitated. The pellet was dissolved in 25 µL of RNase-free water.

#### Ribosomal RNA depletion

Ribosomal RNAs (rRNAs) were removed according to a previous study using NEBNext rRNA Depletion Kit v2 (Human/Mouse/Rat) (New England Biolabs) and a 85 μM DNA oligo mixture containing 85 DNA oligos complementary to either 23S rRNA, 16 S rRNA, or 5 S rRNA. ^115^ 1 µg of the RNA was diluted in RNase-free water for a total volume of 11 µL. The diluted RNA was mixed with 2 µL of NEBNext Probe Hybridization Buffer and 2 µL of the DNA oligo mixture. Annealing was performed by heating the sample for 2 min at 95°C and cooled down to 22°C (0.1°C/sec) in a thermocycler. After additional 5-min incubation at 22°C, rRNA digestion was performed by mixing the sample with 2 µL of RNase H Reaction Buffer, 2 µL of NEBNext Thermostable RNase H, and 1 µL of nuclease-free water, and incubating the sample for 30 min at 50°C in a thermocycler with the lid temperature set to 55°C. The reaction was quickly chilled on ice and mixed with 5 µL of DNase I Reaction Buffer, 2.5 µL of NEBNext RNase-free DNase I, and 22.5 µL of nuclease-free water. This was followed by incubation for 30 min at 37°C in a thermocycler with the lid temperature set to 40°C. The RNA was purified using Monarch RNA Cleanup Kit (New England Biolabs).

#### Library construction

The cleaned-up RNA was used for library construction with NEBNext Ultra II Directional RNA Library Prep Kit for Illumina (New England Biolabs). Reaction was performed according to the manufacturer’s instructions.

#### ChIP-seq

ChIP-seq was carried out as described previously with modifications. ^116^

#### Cell fixation and lysate preparation

For crosslinking, 80 mL of a cell culture were mixed with 2.5 mL of 37% formaldehyde (final concentration: 1%). The mixture was immediately cooled in iced water with occasional agitation. After 5 min, the mixture was incubated for 20 min at room temperature with agitation at 75 rpm. Crosslinking was quenched by adding 4.5 mL of 2.5 M glycine (final 0.13 M). After 20 min of incubation at room temperature, the mixture was transferred to 50-mL tubes while removing as much powder of elemental sulfur as possible. The cells were collected by centrifugation (16,000 x g, 10 min, 4°C) and resuspended in 1 mL of 0.8 × ASW–m1. The cell suspension was transferred to an 1.5- mL tube and centrifuged (10,000 x g, 5 min, 4°C). The pellet was resuspended in 1 mL of TBS-TT (20 mM Tris-HCl [pH 8], 150 mM NaCl, 0.1% v/v Triton X-100, and 0.1% v/v Tween 20) and transferred to a milliTUBE 1 ml AFA Fiber (Covaris) for DNA fragmentation using M220 Focused- ultrasonicator (Covaris). The fragmentation was performed for 12 min at 7°C with the peak power set to 75, the duty factor set to 26, and the cycles/burst set to 200. The lysate was transferred to an 1.5- mL tube and centrifuged (20,400 x g, 30 min, 4°C). The supernatant was transferred to a new 1.5-mL tube, flash frozen in liquid nitrogen, and stored at –80°C until use.

#### Immunoprecipitation

500 μL of the lysate were mixed with 10 µL of antiserum (IP) or TBS-TT and incubated for 2.5 h at 4°C. 50 μL of Dynabeads Protein G (Thermo Fisher Scientific) were washed with TBS-TT and added to the IP sample. The lysate-bead mixture was incubated for 1h at 4°C. The beads were washed five times with TBS-TT, once with TBS-TT containing 0.5 M NaCl, and once with TBS-TT containing 0.5% v/v Triton X-100 and 0.5% v/v Tween 20. Protein-DNA complexes were eluted in 200 μL of elution buffer (20 mM Tris-HCl [pH 7.5], 10 mM EDTA [pH 8], and 0.5% SDS) for 30 min at 65°C with agitation at 1,400 rpm. 186 µL of the input sample were mixed with 4 µL of 0.5 M EDTA (pH 8) and 10 µL of 10% SDS and heated the same way. Each of the IP and input samples was incubated with 3 μL of 800 U/mL proteinase K (New England Biolabs) overnight at 37°C. The DNA was extracted with phenol:chloroform:isoamyl alcohol (Sigma Aldrich) and ethanol-precipitated together with 2 μL of 20 mg/mL glycogen (Thermo Fisher Scientific). The DNA was dissolved in 55 μL of 10 mM Tris-HCl (pH 8.0).

#### Library construction

The DNA was used for library construction with NEBNext Ultra II DNA Library Prep with Sample Purification Beads (New England Biolabs). Reaction was performed according to the manufacturer’s instructions.

### Protein purification and antibody production

#### Purification of XerA

The coding region of *T. kodakarensis* XerA was cloned into the pET21a(+) plasmid (Novagen) and expressed without any additional sequence. *E. coli* cultures were grown to an OD600 of ∼0.8 in LB at 37°C. Protein expression was induced with 0.1 mM IPTG for 15 h at 28°C. The cells were pelleted by centrifugation, resuspended in 1 x PBS, and pelleted again. The cells were resuspended in 50 mM Tris-HCl (pH 7.5) containing 1 mM EDTA and disrupted by sonication. The lysate was cleared by centrifugation and heated for 20 min at 70°C. The heat-denatured proteins were removed by centrifugation. The supernatant was concentrated using an Amicon Ultra Centrifugal Filter 10 kDa MWCO (Merck) and loaded onto a HiTrap Heparin HP affinity column (Cytiva) and eluted with a linear gradient of NaCl (0 to 1 M). The main peak fractions were pooled and fractionated through a Superdex-200 10/300 GL column (Cytiva) using 50 mM Tris-HCl (pH 7.5) containing 150 mM NaCl as a mobile phase. The main peak fractions were pooled and concentrated using an Amicon Ultra Centrifugal Filter 10 kDa MWCO (Merck). The sample was stored in the presence of 650-800 mM NaCl to avoid precipitation.

#### Purification of Smc

The coding sequence of *T. kodakarensis* Smc with a 5’ extension of ATGTGG was cloned into pET21a(+) and expressed without any additional sequence. Cells harboring the expression plasmid were cultured in LB medium containing 50 μg/mL of ampicillin and 34 μg/mL of chloramphenicol. The culture was grown to an OD600 of ∼0.4 in LB at 37°C. Expression of the *smc* gene was induced with 1 mM IPTG for 4 h at 37°C. The cells were pelleted by centrifugation and resuspended in 50 mM Tris-HCl (pH 8) containing 1 mM EDTA and 500 mM NaCl. After cell disruption by sonication, the cell extract was obtained by centrifugation followed by heating for 20 min at 80°C. The heat- denatured proteins were removed by centrifugation. The supernatant was mixed with polyethylenimine (final 0.15%) and incubated for 10 min on ice. Precipitated nucleic acids were removed by centrifugation (23,708 x g, 10 min, 4°C). Ammonium sulfate was added to the supernatant to 70% saturation, and the soluble proteins were precipitated by overnight incubation at 4°C. The proteins were recovered by centrifugation (23,708 x g, 20 min, 4°C). The precipitate was dissolved in 50 mM Tris-HCl (pH 8) containing 1 mM EDTA and 1 M ammonium sulfate. Debris were removed from the solution through a 0.45-μm Minisart Filter (Sartorius). The sample was loaded onto a HiTrap Phenyl HP column (Cytiva), and the chromatography was developed with a linear gradient of ammonium sulfate (1 to 0 M). The main peak fractions were pooled and dialysed overnight against 50 mM Tris-HCl (pH 8) containing 1 mM EDTA and 100 mM NaCl. The dialysed sample was loaded onto a HiTrap Heparin HP affinity column (Cytiva). Elution was performed with a linear gradient of NaCl (0 to 1 M). The main peak fractions were pooled and diluted in 50 mM Tris-HCl (pH 8) containing 1 mM EDTA so that the NaCl concentration was adjusted to 100 mM. Lastly, the Smc protein was purified through a HiTrap Q HP column (Cytiva) with a linear gradient of NaCl (0.1 to 1 M).

#### Purification of TrmBL2

The coding sequence of *T. kodakarensis* TrmBL2 was cloned into pET21(+) and expressed without any additional sequence. The expression plasmid was kindly provided by Hugo Maruyama. ^55^ Protein expression was induced as described previously^55^. A heat-stable protein extract was prepared and concentrated as described for XerA purification, except that heating was performed at 80°C. It was loaded onto a HiTrap Heparin HP affinity column (Cytiva) and eluted with a linear gradient of NaCl (0 to 2 M). A pool of the main peak fractions were loaded onto a ResourceQ anion exchange column (Cytiva) and eluted with a linear gradient of NaCl (0 to 1 M). The main peak fractions were pooled and concentrated using an Amicon Ultra Centrifugal Filter 10 kDa MWCO (Merck). The sample was stored in 50 mM Tris-HCl (pH 8.0) containing 240 mM NaCl.

#### Antibody production

The purified recombinant proteins were used to raise rabbit polyclonal antibodies (Eurofins Genomics).

### DNA affinity purification

#### DNA probes

PCR was performed to amplify *difL* and *difR* (using genomic DNA of *T. kodakarensis* as a template) and *kanR* (using pGBKT7 [Takara Clontech] as a template) and clone them separately into the pUC19 plasmid (digested at the BamHI and SphI sites) using In-Fusion HD Cloning Plus reagents. The primer sets used for the cloning are shown in Table S1. The inserted fragments were PCR amplified to generate biotinylated DNA probes (see Table S1 for oligonucleotide sequences). The probes were purified using FastGene Gel/PCR Extraction Kit (Nippon Genetics).

#### Cell extract preparation and affinity purification

80 mL of a cell culture were dispensed into two 50-mL tubes, chilled in iced water for 10 min, and centrifuged (10,000 x g, 10 min, 4°C). The two pellets were resuspended together in 3.5 mL of buffer H (20 mM Tris-HCl [pH 7.5]、500 mM NaCl) and sonicated. The lysate was cleared by centrifugation (12,000 x g、10 min, 4°C). 3 mL of the supernatant were mixed with 7 mL of 20 mM Tris-HCl (pH 7.5) containing 0.714% v/v Tween-20. The diluted lysate corresponding to 0.9 mg protein was mixed with 45 µL of 2 mg/mL Poly(dI-dC) (Sigma Aldrich). The sample was mixed with Dynabeads MyOne Streptavidin T1 (Thermo Fisher Scientific) pre-coupled to 9 µg of a biotinylated probe. The mixture was incubated for 30 min at 5°C with agitation at 600 rpm. The beads were recovered and resuspended in 1.5 mL of buffer MT (20 mM Tris-HCl [pH 7.5], 150 mM NaCl, 0.5% v/v Tween-20). Beads were recovered from 500 µL of the sample for total bound proteins and from 1,000 µL for eluates 1 and 2 in Figure 5.

The beads for total bound proteins were washed with buffer MT and resuspended in 30 µL of buffer W600 (20 mM Tris-HCl [pH 7.5], 600 mM NaCl, 0.5% v/v Tween-20). The sample was mixed with 15 µL of 3 x Blue Protein Loading Dye (New England Biolabs) and incubated for 10 min at 37°C with agitation at 600 rpm to elute the bound proteins. The supernatant was recovered, mixed with 1.5 µL of 1.25 M DTT, and heated for 5 min at 95°C.

The beads for eluates 1 and 2 were first washed with buffer MT. Proteins were then sequentially eluted from the beads with (1) 56.2 µL of buffer W300 (20 mM Tris-HCl [pH 7.5], (2) 300 mM NaCl, 0.5% v/v Tween-20), (3) 58.1 µL of buffer W450 (20 mM Tris-HCl [pH 7.5], 450 mM NaCl, 0.5% v/v Tween-20), (4) 60 µL of buffer W600, (5) 48 µL of buffer W750 (20 mM Tris-HCl [pH 7.5], 750 mM NaCl, 0.5% v/v Tween-20), (6) 36 µL of buffer W1000 (20 mM Tris-HCl [pH 7.5], 1 M NaCl, 0.5% v/v Tween-20), and (7) 90 µL of SDS sample buffer (buffer W600 containing 1 x Blue Protein Loading Dye). Each elution was performed by incubating the bead suspension for 10 min at 37°C with agitation at 600 rpm and recovering the supernatant. The supernatant from (5), used as eluate 1, was mixed with 12 uL of 20 mM Tris-HCl (pH 7.5) and 30 µL of 3 x Blue Protein Loading Dye. The supernatant from (7) was used as eluate 2. Eluates 1 and 2 were each mixed with 3 µL of 1.25 M DTT and heated for 5 min at 95°C.

#### Detection of bound proteins

20 µL of the DNA affinity purification sample per lane were subjected to SDS-PAGE using e-PAGEL HR 5-20% gradient gel (ATTO). The gel was stained with Pierce Silver Stain for Mass Spectrometry (Thermo Fisher Scientific). Indicated gel bands were excised, digested with trypsin, and analyzed by NanoLC-MS/MS using a Q Exactive mass spectrometer (Thermo Fisher Scientific) and an Ultimate 3000 nanoLC pump (AMR). Peptides and proteins were identified by means of automated database searching using Sequest HT (Thermo Fisher Scientific) against the *T. kodakaraensis* database (Taxon ID 311400, UniProtKB/SWISS-PROT, release 2018-01) with a precursor mass tolerance of 10 p.p.m., a fragment ion mass tolerance of 0.02 Da, and trypsin specificity that allows for up to two missed cleavages. A reversed decoy database search was conducted with Percolator node, setting the false discovery rate (FDR) threshold to 5% at the peptide level.

### Identification of putative *dif* sequences near domain boundaries

Prior work reported nine Smc-dependent boundaries and their closest genes in *H. volcanii*. ^38^ We first retrieved the sequences of these genes together with their upstream and downstream 4-kb regions. To identify potential boundary-associated *dif* sequences, we searched these nine regions for imperfect inverted repeats containing TAA(N)nTTA (n = 6-8). We selected this motif since it is the most highly conserved feature among the previously reported *dif* sequences in archaea (Figure S4A). We identified three sequences containing the motif, one of which is located upstream of the *xerA*-containing operon as illustrated in Figure 4E. Another hit is located upstream of *HVO_B0249* as shown in Figure S4C. The third hit (AATC<u>G</u>AG<u>AGTAA</u>GGAGAGTA<u>TTACT</u>AA<u>C</u>CGGC, matches between the imperfect repeats are underlined) is located ∼200 bp away from the second one. The third sequence was removed from the candidates due to a smaller number of base matches than that in the second one. The *dif*-like sequence in *H. salinarum* was identified by searching for the intergenic region upstream of the same *xerA*-containing operon.

## QUANTIFICATION AND STATISTICAL ANALYSIS

### Illumina DNA sequencing and read mapping

The 3C-seq, RNA-seq, and ChIP-seq libraries generated in this study were paired-end sequenced on either of the Illumina HiSeq X Ten, NovaSeq 6000, NextSeq 550, and NextSeq 500 platforms. The sequencing was performed by Macrogen, Single-cell Genome Information Analysis Core at Kyoto University, and the NGS core facility of the Graduate Schools of Biostudies at Kyoto University. Sequence data were mapped to the reference genome of *T. kodakarensis* KOD1 (GCA_000009965.1), *T. acidophilum* DSM 1728 (GCA_000195915.1), *H. volcanii* DS2 (GCA_000025685.1) or *H. salinarum* NRC-1 (GCA_000006805.1). The accession numbers of the sequence data from prior work^38^ are as follows. *T. kodakarensis*: SRR12717836 and SRR12717848, *H. volcanii* DS2: SRR11747722 and SRR11747734, *H. volcanii* Δ*smc*: SRR11747720 and SRR11747731, *H. salinarum*: SRR12717837 and SRR12717838.

### 3C-seq data analysis

#### Generation of 3C-seq contact maps

To generate 3C-seq contact maps, reads from two to three replicates were combined and processed using HiC-Pro (version 3.0.0). ^98^ The data were iteratively corrected with the MAX_ITER parameter of 500. The obtained matrices were normalized so that the sum of interaction scores is equal to 1000 for each row and column. For *T. kodakarensis*, genomic bins with extremely low coverage were filtered out by setting the FILTER_LOW_COUNT_PER parameter to 0.006. 3C-seq contact maps reflecting the genomic inversion in *T. kodakarensis* were generated by flipping the sequence corresponding to the genomic coordinates 327001-520000.

#### Insulation score analysis

Insulation score was determined as described previously at 1-kb resolution. ^36^ The size of the sliding square was set to 40 kb.

#### Detection of loop structures

DNA loops in the *T. kodakarensis* strain KU216 were searched for using Chromosight (version 1.5.0) ^48^ as described previously. ^36^ 3C-seq contact maps at 1-kb resolution were used for the analysis. A genomic segment containing the inversion (the genomic coordinates 321001-525000) were omitted due to the uncertainty resulting from the genome heterogeneity caused by the inversion. The max- dist parameter was set to 1044000.

### RNA-seq data analysis

Differential gene expression analysis was performed using RNA-seq data from three replicates and the GFF annotation file of the *T. kodakarensis* reference genome. The 23S, 16S, and 5S rRNA genes were omitted from the analysis. The orientation of the 7S (SRP) RNA was inverted, because the original orientation in the genome annotation is probably incorrect as reported previously. ^117^ RNA levels were first quantified by Salmon (version 1.9.0) ^99^ and then analyzed using edgeR (version 3.40.2) ^100^ as follows. The read count was normalized using the TMM normalization method. Common, trended, and tagwise dispersions were estimated using the GLM method. Statistical significance of the gene expression difference was tested using the glmFit function.

### ChIP-seq data analysis

#### Mapping

Reads from two replicates were combined and mapped using Bowtie 2 (version 2.3.5.1). ^101^ Low- quality alignments (MAPQ < 30) were removed using SAMtools (version 1.9). ^102^

#### Generation of ChIP-seq tracks

Generated BAM files were processed using the bamCoverage function (version 3.5.1) of deepTools^103^ to calculate Reads Per Kilobase region per Million mapped reads (RPKM) for genomic bins of 50 bp and 1 kb. RPKM ratios of immunoprecipitated versus input DNA were plotted as IP/input.

#### Peak analysis

Mappled data were processed for peak calling using MACS (version 3.0.0a6) ^104^ with the following setting: --fe-cutoff 3, --keep-dup all, --call-summits, -m 1 50. To identify common binding sites for Smc and TrmBL2, overlaps of ±100-bp regions from the identified peaks were examined using the numOverlaps function of regioneR (version 1.30.0). ^105^ Statistical significance of the overlaps was evaluated using the overlapPermuTest function of regioneR. DNA motifs enriched at identified peaks were determined using MEME-ChIP (version 5.5.5) ^97^ with the following setting: -ccut 100, -order 2, -minw 4, -maxw 15, -meme-mod zoops. To calculate protein occupancy at the identified peak, the read coverage of the ±100-bp region from the peak summit was calculated using the multicov function of BEDtools (version 2.26.0). ^106^ The coverage was normalized to the total read number, and the IP/input ratio of the normalized coverage values was used as the occupancy of the protein at the peak.

#### Smc stalling efficiency

Using ChIP-seq data from KU216, we first calculated Smc occupancy for the Smc peaks overlapping with those of TrmBL2. Smc occupancy in Δ*trmBL2* was also calculated for the same regions. These values were used to calculate the Smc occupancy ratio between KU216 and Δ*trmBL2*. Loci with the Δ*trmBL2*/KU216 ratio of –0.5 or larger were filtered out as less probable stalling sites. We then calculated TrmBL2 occupancy for the TrmBL2 peaks overlapping with the retained Smc peaks in KU216. These peak pairs were used to calculate the ratio of Smc versus TrmBL2 occupancies, which was defined as Smc stalling efficiency.

### Analysis on DNA sequence features

AT-content tracks were generated using SeqKit (version 2.5.1) ^107^ with sliding window size of 200 bp and step size of 100 bp. For analysis on stalling sites, 200-bp sequences centered at TrmBL2 peaks were used. Occurrence of DNA tetranucleotide sequences was counted for each stalling site using the countDnaKmers function of seqTools (version 1.32.0). For non-palindrome sequences, their occurrence frequencies plus those of the reverse complements were used for analyses. The occurrence frequency dataset was then used to identify the tetranucleotide sequences that were correlated with Smc stalling efficiency. The same dataset was used to calculate the average of 4-bp-scale persistence lengths for each stalling site. The persistence length values were according to a previous study. ^69^ To calculate DNA cyclizability, we predicted normalized cyclizability scores (*C*-scores) of the 50-bp bins along each sequence using the DNAcycP web server (https://dnacycp.stats.northwestern.edu/).^70^ The average of the *C*-scores was calculated for each stalling site and used as its cyclizability score for correlation analysis. The other DNA properties were predicted using the Deep DNAshape web server (https://deepdnashape.usc.edu/)^72^ with the Deep DNAshape layer set to 4. This serve predicts a selected feature value for each base of the input sequence. The base average of the feature value was calculated for each stalling site and used for correlation analysis. Adjusted *p*-values were determined using the Benjamini-Hochberg method.

### Coarse-grained MD simulations of stalling sites

The coarse-grained MD simulations were conducted with CafeMol (version 3.2.1). ^108^ The 3SPN.2C sequence-dependent coarse-grained DNA model was used. ^79^ 5’- and 3’ termini of stalling sites were capped with five CG repeats to insulate the stalling sites from end effects, but the CG caps were omitted when gross dynamic PLs were calculated from the obtained MD data. The temperature was set to 300 K. The monovalent ion concentration of Debye-Hückel model was set to 150 mM. Langevin dynamics simulations were performed with a step size of 0.2 in the CafeMol time unit for 2×10^8^ steps per stalling site, including 5×10^7^ steps for equilibration. The coordinates were stored every 5×10^3^ steps.

The gross dynamic PL, *l*_*d*_, was estimated from the directional correlation decay of DNA polymer as 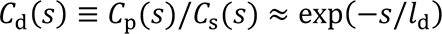 as performed previously. ^68^ Here, *C*_*s*_ is the autocorrelation function of the helical axis vectors *h* defined as 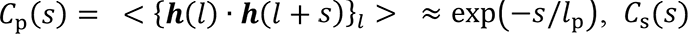 is that for minimum-energy ground state conformation defined as 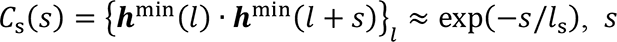 is the spacing between helical axis vectors, *l* is the position along the contour length, *l*_*p*_ is the PL, and *l*_*s*_ is the static PL. *s* is a multiple of 10 in order to remove any residual effects of DNA helicity. {η} represents an average over the position along the contour length and <η> represents a long-time average. The sequence-dependent ground state conformation was obtained using x3DNA (version 2.4). ^109^

### Data visualization and statistical test

Unless otherwise stated, data visualization and statistical test was performed using R software (http://www.r-project.org/).

## SUPPLEMENTARY FIGURES

**Figure S1.**
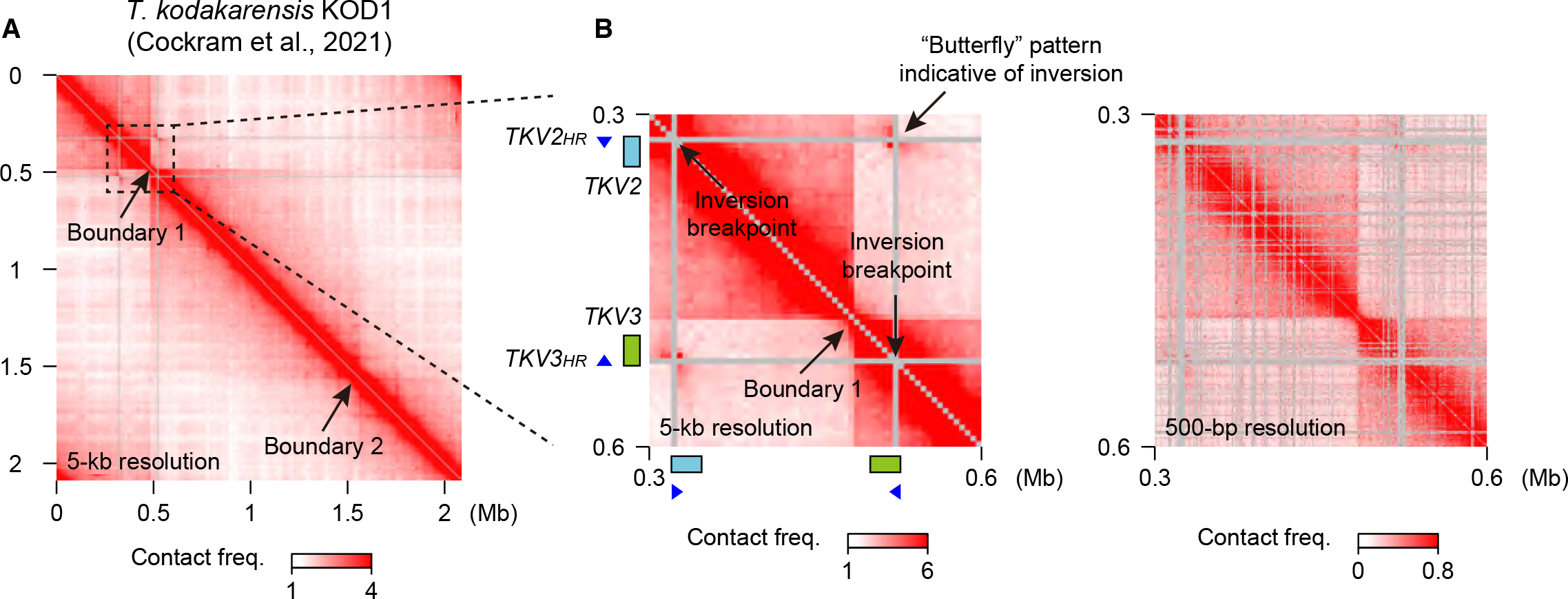
Analysis on published Hi-C data of the wild-type *Thermococcus kodakarensis* strain KOD1. Related to Figure 1. (A) A Hi-C contact map of the KOD1 genome was generated as in Figure 1A using previously published Hi-C data. ^38^ (B) Magnified contact maps are shown as in Figure 1B at 5-kb (left panel) and 500-bp (right panel) resolutions.

**Figure S2.**
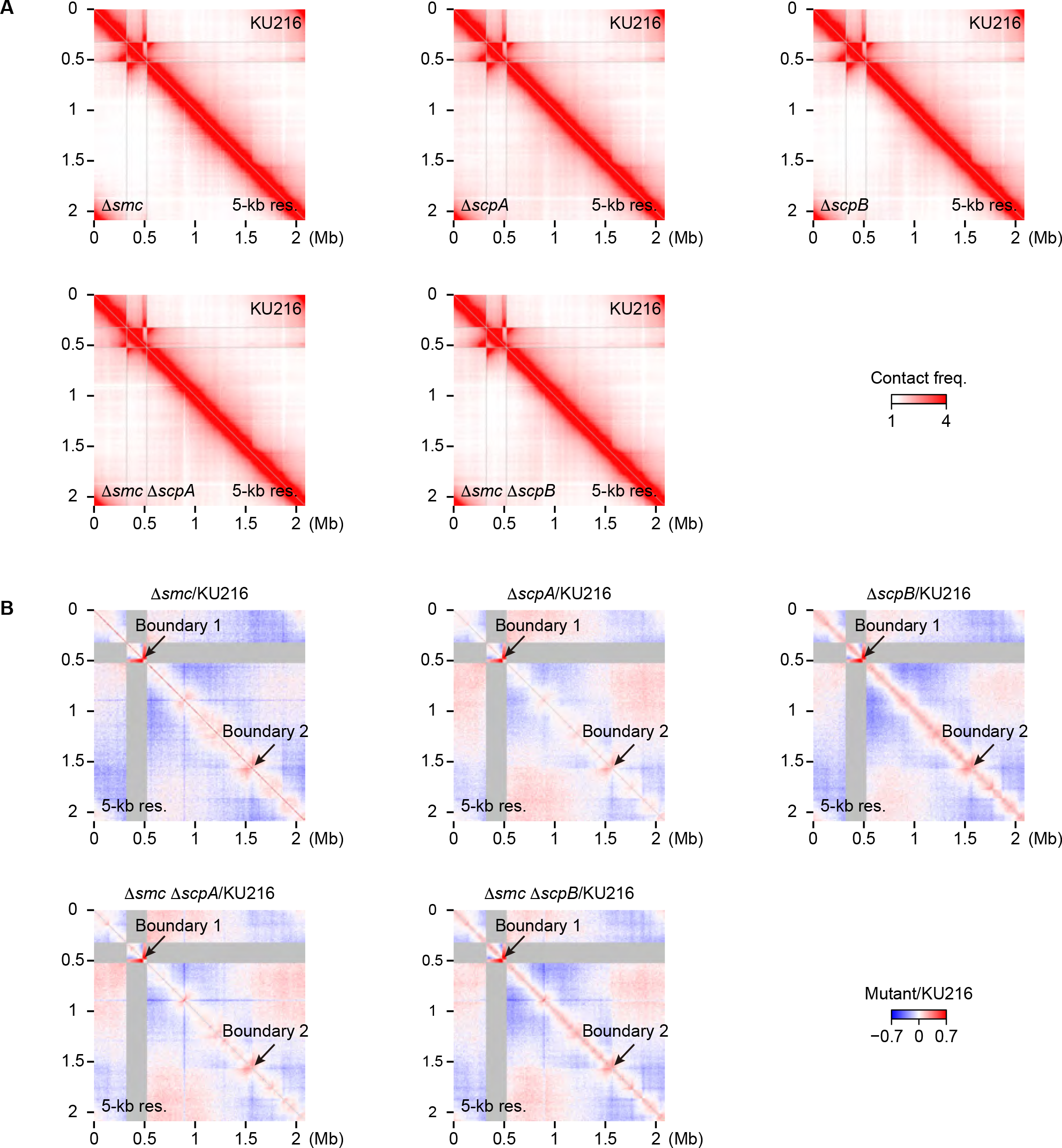
Deletions of *smc*, *scpA*, and *scpB* have similar impacts on genome organization. Related to Figure 2. (A and B) Comparison of 3C-seq contact maps (5-kb resolution) from KU216 and deletion strains (upper right triangles and lower left triangles, respectively). (B) Differential contact maps showing log2 ratios of contact frequencies between KU216 and deletion strains were generated at 5-kb resolution. Locations of boundaries 1 and 2 in KU216 are indicated by arrows. The genomic contacts between the inverted region and other loci are omitted from the analysis.

**Figure S3.**
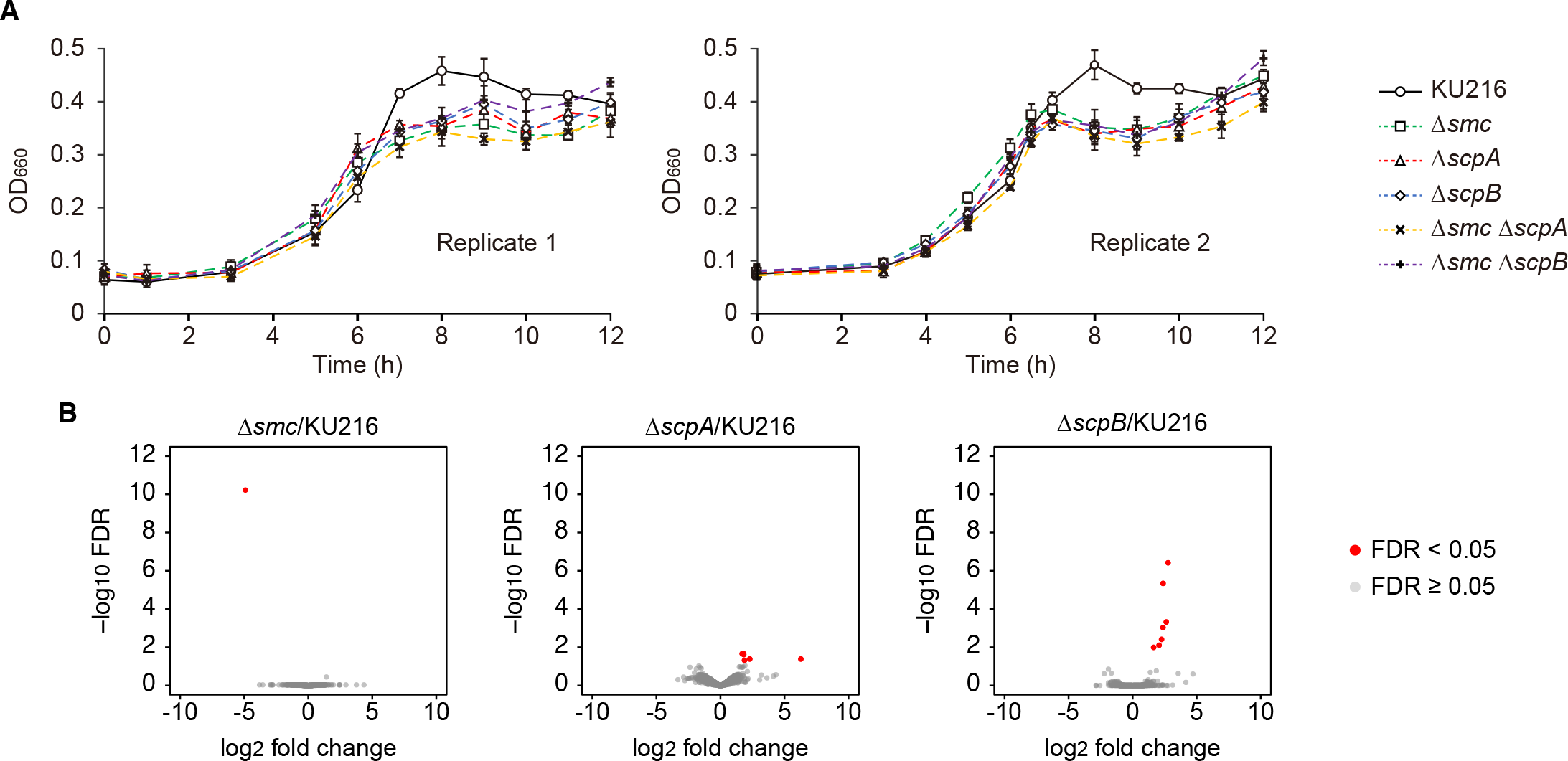
Phenotypic analysis on Smc-ScpAB deletion strains. Related to Figure 2. (A) Growth curves of KU216 and deletion strains in nutrient-rich medium (ASW-YT-m1-S^0^). Error bars represent standard deviations from triplicate cultures grown in each replicate. (B) Volcano plots showing gene expression changes and their significance in Δ*smc* (left), Δ*scpA* (middle), and Δ*scpB* (right) versus KU216. Differentially expressed genes with false discovery rate (FDR) values below 0.05 are shown in red.

**Figure S4.**
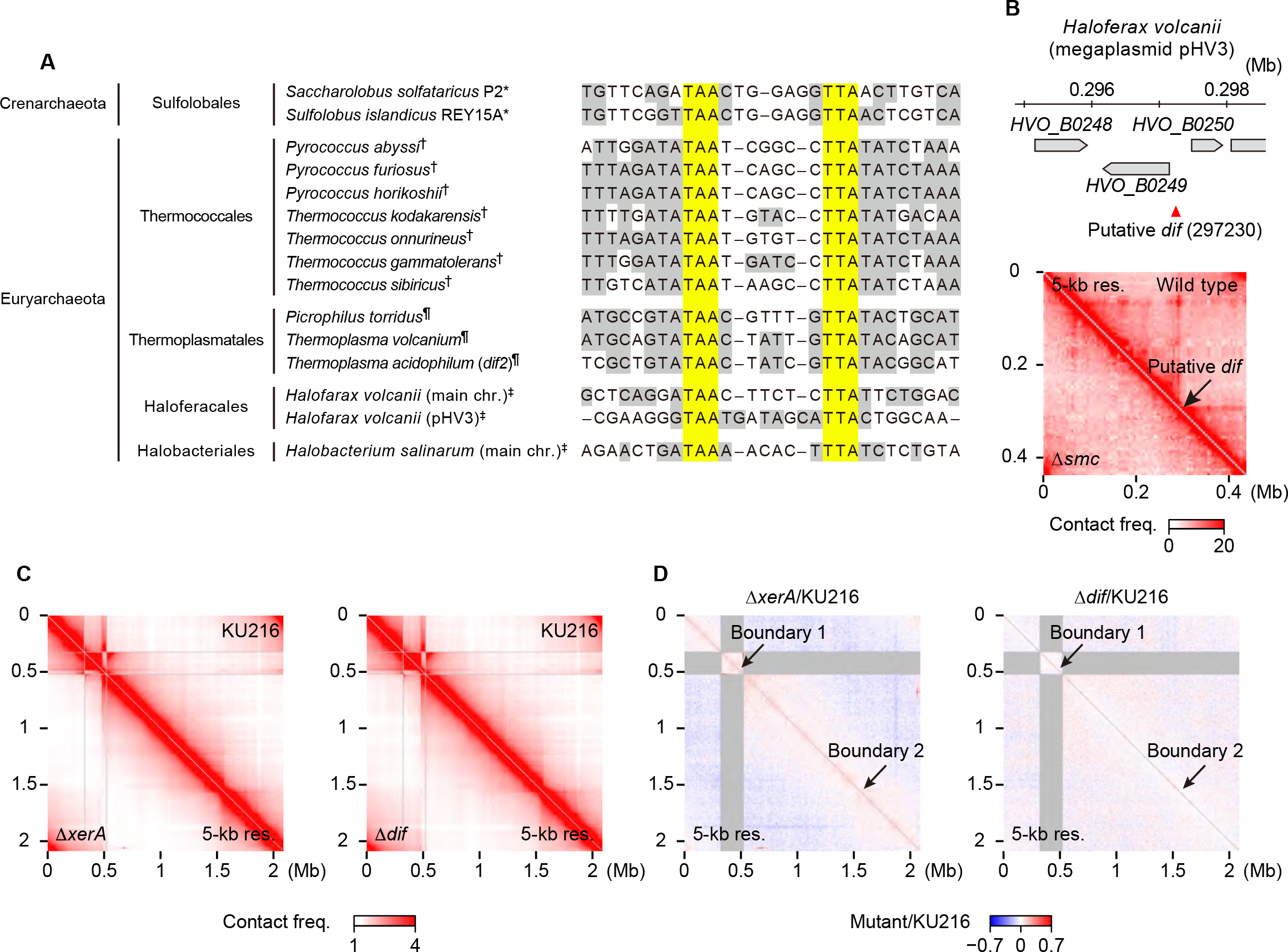
Characterization of the archaeal Xer/*dif* system. Related to Figures 3 and 4. (A) Multiple sequence alignment of archaeal *dif* sequences. Shading indicates matching bases between the incomplete inverted repeats in each sequence. Yellow shading highlights matching bases common to all sequences. *Duggin et al.. ^51^ ^†^Cossu et al.. ^118^ Jo et al.. ^53^ ^
‡^ Putative *dif* sequences identified in this study. (B) Upper panel: the genomic position of a putative *dif* sequence on the megaplasmid pHV3 in *Haloferax volcanii*. Coordinates of the first bases of the *dif* sequences are in parentheses. Neighboring genes and their orientations are indicated by gray and pentagons. Lower panel: published Hi-C data^38^ were used to generate contact maps (5-kb resolution) of pHV3 in wild-type and Δ*smc* cells of *H. volcanii* (upper right and lower left triangles, respectively). The position of the putative *dif* sequence is indicated by an arrow. (C) Comparison of 3C-seq contact maps (5-kb resolution) from KU216 and deletion strains (upper right triangles and lower left triangles, respectively). (D) Differential contact maps displaying log2 ratios of contact frequencies between KU216 and deletion strains are shown as in Figure S2B.

**Figure S5.**
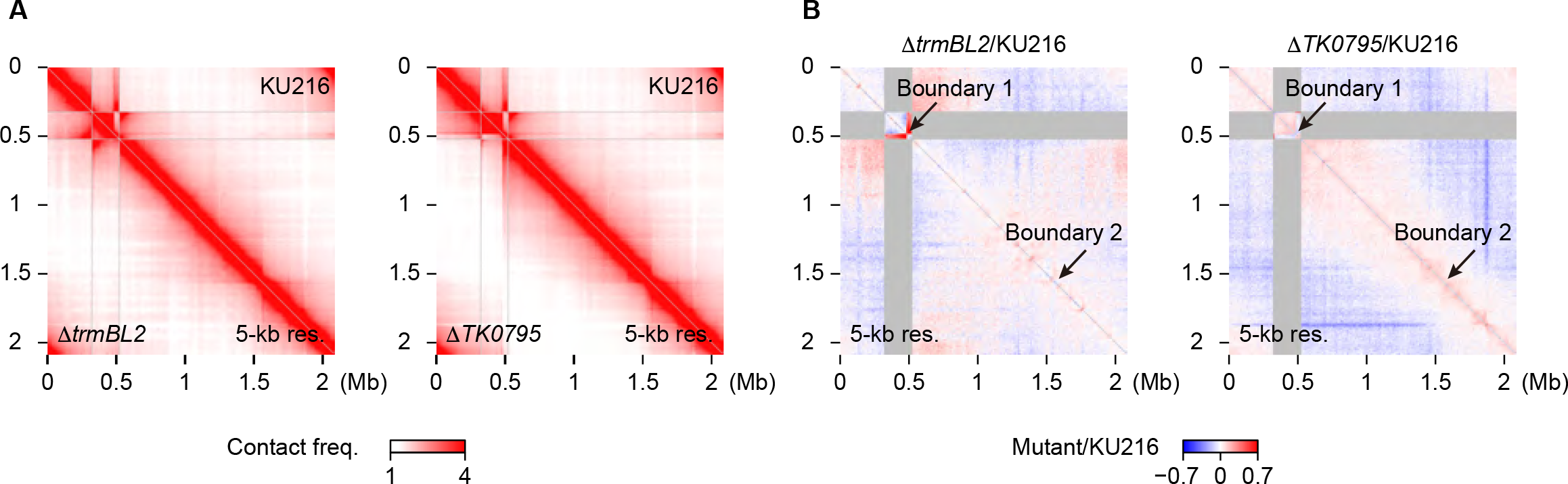
Effects of *trmBL2* and *TK0795* deletions on the 3D genome. Related to Figure 5. (A) Comparison of 3C-seq contact maps (5-kb resolution) from KU216 and deletion strains (upper right and lower left triangles, respectively). (B) Differential contact maps displaying log2 ratios of contact frequencies between KU216 and deletion strains are shown as in Figure S2B.

**Figure S6.**
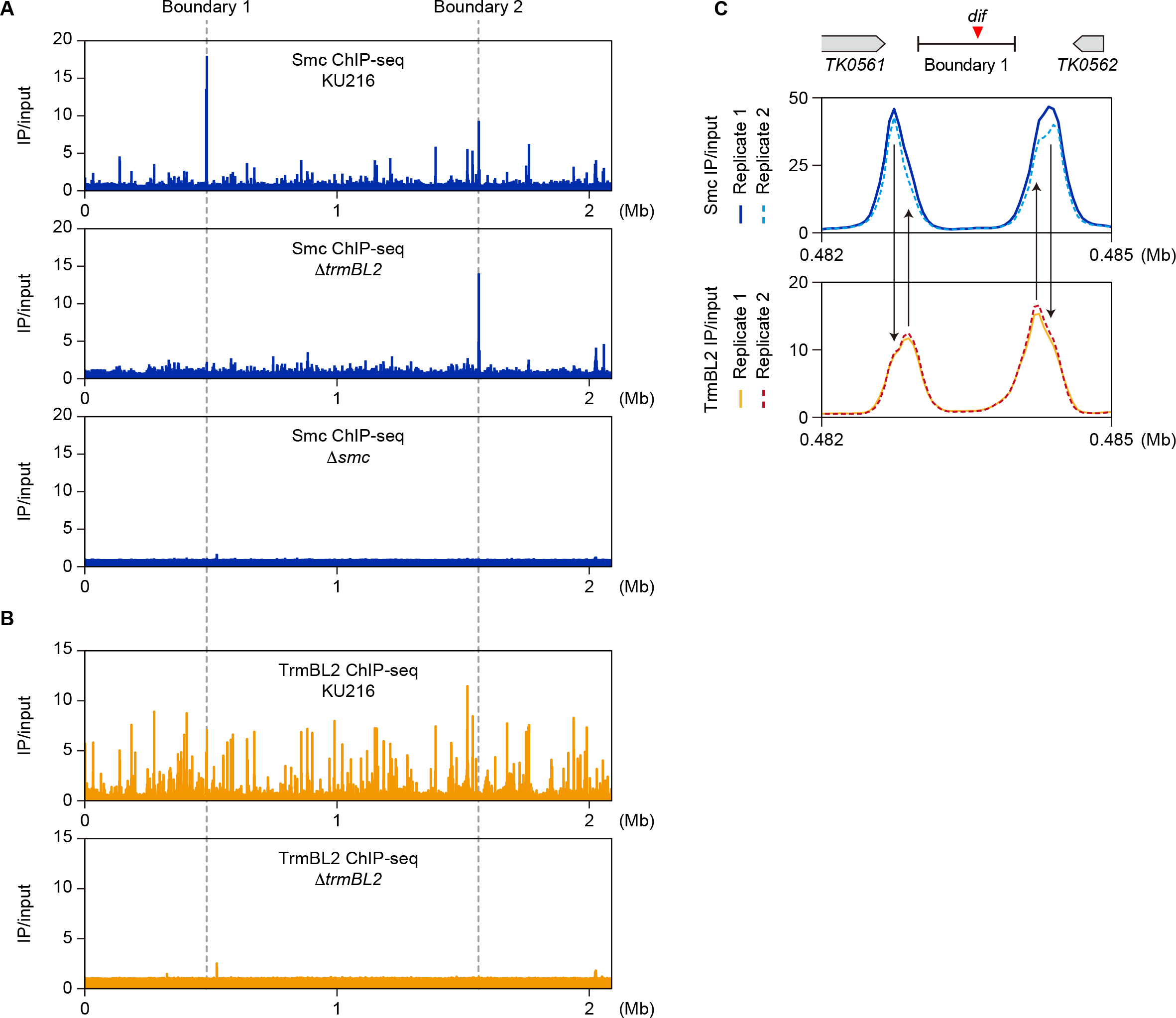
ChIP-seq analysis of Smc and TrmBL2. Related to Figure 6. (A) ChIP-seq tracks of Smc for the whole genomic regions of KU216, Δ*trmBL2*, and Δ*smc* strains. Enrichment of immunoprecipitated versus input DNA (IP/input) is shown at 1-kb resolution. The positions of boundaries 1 and 2 are indicated by gray dotted lines. (B) ChIP-seq tracks of TrmBL2 for the whole genomic regions of KU216 and Δ*trmBL2* strains are shown as in (A). (C) ChIP-seq tracks of Smc and TrmBL2 at boundary 1 are shown for two replicates obtained from KU216. Data are shown as in Figure 6A. The differences in the peak summit positions of Smc and TrmBL2 are highlighted by arrows.

**Figure S7.**
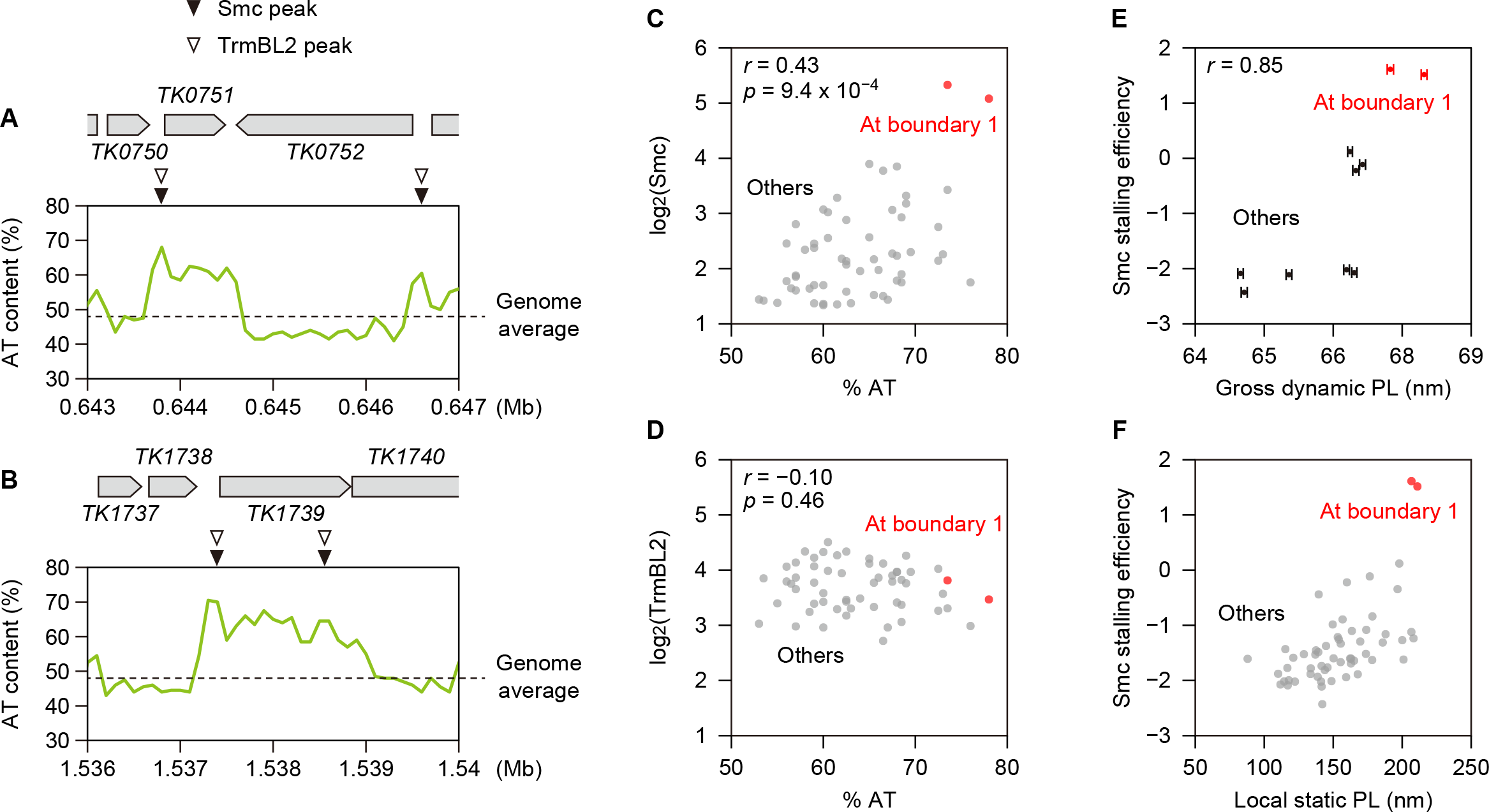
AT-rich sequences are associated with Smc-ScpAB stalling. Related to Figure 7. (A and B) AT-content tracks of the non-boundary loci shown in Figures 6C and D. Data are shown as in Figure 7A. (C) AT content and TrmBL2 occupancy were plotted for 57 stalling sites. The Spearman rank correlation coefficient (*r*) and corresponding *p*-value are also shown. The two stalling sites associated with boundary 1 are highlighted in red. (D) AT content and Smc occupancy were plotted for the 57 stalling sites. The Spearman rank correlation coefficient (*r*) and corresponding *p*-value are also shown. The two stalling sites associated with boundary 1 are highlighted in red. (E) MD-based gross dynamic persistence length (PL) and Smc stalling efficiency were plotted for the 5 most efficient and 5 most inefficient stalling sites among the 57 sites. The Spearman rank correlation coefficient (*r*) is also shown. Errors of the gross dynamic PLs were estimated using the jackknife method. The two stalling sites associated with boundary 1 are highlighted in red. (F) Local static PL and Smc stalling efficiency were plotted for the 57 stalling sites. The two stalling sites associated with boundary 1 are highlighted in red.

## Notes

### Competing Interest Statement

The authors have declared no competing interest.

